# Development of semisynthetic blasticidin S analogs with potent and fast-killing anti-malarial activity

**DOI:** 10.64898/2026.04.12.718010

**Authors:** Katherine R. Fike, Cole Gannett, Anastazja M. Kiselka, Kateland Tiller, Toheeb Ajasa, James Weger-Lucarelli, Anne M. Brown, Andrew N. Lowell, Michael Klemba

**Affiliations:** Department of Biochemistry, Virginia Tech, Blacksburg, VA 24061, United States; Center for Emerging, Zoonotic, and Arthropod-borne Pathogens, Virginia Tech, Blacksburg, VA 24061, United States; Virginia Tech Center for Drug Discovery, Virginia Tech, Blacksburg, VA 24061, United States; Department of Chemistry, Virginia Tech, Blacksburg, VA 24061, United States; Department of Biomedical Sciences and Pathobiology, Virginia-Maryland College of Veterinary Medicine, Virginia Tech, Blacksburg, VA 24061, United States; Center for One Health Research, Virginia Tech, Blacksburg, VA 24061, United States; Research and Informatics, University Libraries, Virginia Tech, Blacksburg, VA 24061, United States; Faculty of Health Sciences, Virginia Tech, Blacksburg, VA 24061, United States

**Keywords:** Plasmodium, malaria, blasticidin S, ribosome, inhibitor

## Abstract

Protein synthesis represents an attractive target space for the development of anti-malarials with novel modes of action. Natural-product inhibitors of the eukaryotic 80S ribosome can have potent anti-malarial activity but are often poorly selective due to mammalian cytotoxicity. Blasticidin S (BlaS) is a microbially-produced natural product that broadly inhibits prokaryotic and eukaryotic protein synthesis by binding to the ribosomal peptidyl transferase center. In this study, we explored the potential for improving the anti-malarial potency and selectivity of the blasticidin S scaffold with semi-synthetic analogs that are modified at the C6’ and C4 sites. The two best analogs were two orders of magnitude more potent than BlaS against *Plasmodium falciparum* drug-sensitive and -resistant lines while displaying low cytotoxicity towards mammalian cells. These analogs exhibited improved kinetics of inhibition of protein synthesis in cultured parasites and blocked the development of asexual stages expressing the plasmodial surface anion channel, a transporter required for nutrient acquisition and BlaS uptake. They also exhibited a dramatically improved speed of killing over BlaS. Molecular docking analysis revealed that these analogs are able to form more interactions with the *P. falciparum* ribosomal peptidyl transferase center than is BlaS, which is consistent with their increased potency. Together, these studies demonstrate the feasibility of generating BlaS analogs with potent anti-malarial activity and provide a roadmap for further development.

---

Over the past 25 years, substantial progress has been achieved in reducing morbidity and mortality caused by malaria. Spurred by control modalities such as the insecticide-treated bed nets and artemisinin combination therapy (ACT), malaria-associated mortality declined substantially from 2000 to 2015.^1^ However, progress has plateaued since 2020, with the number of annual deaths remaining at around 600,000.^1^ Alarmingly, localized high rates of treatment failure with several ACTs have been observed,^1^ potentially threatening these valuable therapeutic resources. To prepare for the eventual widespread clinical failure of ACTs, anti-malarial therapies with new modes of action are urgently needed.

Parasite protein synthesis has emerged as an attractive target space for anti-malarial development because robust translation is required during the pathogenic, asexual replication cycle within host erythrocytes. For *Plasmodium falciparum*, this 48-hour cycle begins with merozoite invasion and culminates in the production of 16-24 daughter parasites, placing a high, sustained demand on translational capacity. Translation occurs in three distinct locations: the cytosol and two parasite organelles, the mitochondrion and the apicoplast. The apicoplast ribosome is the target of numerous antibiotics that inhibit prokaryotic translation, (*e.g.*, doxycycline and azithromycin). These agents elicit a “delayed death” phenotype in which parasite death occurs during the subsequent replication cycle,^2, 3^ making them relatively slow-acting and therefore less desirable candidates for treatment of clinical malaria. In contrast, inhibition of the cytosolic translation apparatus can suppress parasite growth more rapidly, motivating efforts to identify potent, selective inhibitors of this promising target.^4–6^ No inhibitors targeting the mitochondrial ribosome have been reported.

Given its central role in translation, the malarial cytosolic 80S ribosome is an obvious target for translation-directed inhibitors. Many ribosome-inhibiting natural products bind to the peptidyl transferase center (PTC), which is comprised of the exit (E-), peptidyl (P-), and aminoacyl (A-) sites. Potent anti-malarial efficacy has been demonstrated for numerous compounds that target the E- and A-sites of the eukaryotic ribosome and inhibit elongation, for example cycloheximide (E-site) and anisomycin (A-site).^5^ Emetine is an ipecac alkaloid historically used as an anti-amoebic drug but abandoned because of dose-limiting cardiotoxicity and myopathy.^7^ Emetine exerts potent anti-malarial activity by binding to the E-site of the *P. falciparum* ribosome^8, 9^ and its safer analog dehydroemetine has renewed interest in this scaffold for anti-malarial development.^10^ The natural product pactamycin and its analog 7-deoxypactamycin are broad-spectrum E-site inhibitors with extremely potent anti-malarial activities and high cytotoxicity.^11, 12^ Further development of pactamycin analogs that exhibit improved selectivity for the parasite has been reported, which suggests that selective targeting of the malarial ribosome may be feasible despite the overall high conservation of the eukaryotic 80S PTC.^13, 14^ The anti-malarial drug mefloquine was proposed to inhibit the *P. falciparum* cytosolic ribosome by binding to the GTPase-associated center;^15^ however, subsequent work using a *P. falciparum in vitro* translation assay found no translation-inhibitory activity for mefloquine,^5^ suggesting a primary mechanism other than direct inhibition of the ribosome. These examples highlight cytosolic translation, and specifically the 80S ribosome, as a promising, validated target, but also underline the challenge of establishing parasite selectivity.

Blasticidin S (BlaS, **1**, **Fig. 1**) is an antimicrobial, natural product ribosome inhibitor, but its cross-kingdom activity and associated cytotoxicity have precluded clinical development; instead, it has found widespread use as a selectable marker in eukaryotic systems, including *Plasmodium* spp.^16^ Structural studies of eukaryotic ribosomes have revealed that BlaS occupies the P-site and interacts exclusively with the 16S rRNA, occluding the binding of an aminoacyl-tRNA to the A-site.^17, 18^ In prokaryotes, BlaS mainly inhibits termination, whereas in mammalian cells it acts primarily as an elongation inhibitor.^18, 19^ BlaS is highly polar and entry into infected erythrocytes is dependent on a “plasmodial surface anion channel” (PSAC), which normally mediates uptake of essential nutrients.^20^ Accordingly, modest BlaS resistance can be conferred by reducing PSAC-mediated transport, but this incurs a substantial fitness cost.^21, 22^

**Figure 1.**
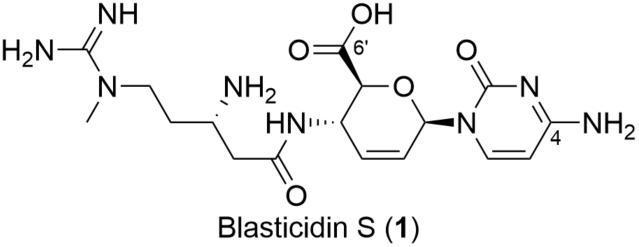
The natural product blasticidin S.

We have recently demonstrated that BlaS can be semi-synthetically modified to improve antimicrobial activity while reducing mammalian cytotoxicity.^23–25^ Encouraged by these findings, we asked if BlaS could be re-engineered into a potent antimalarial with improved host-cell tolerability. Here, we explore two complementary BlaS modification pathways: amidation of the C6’ carboxylate and acylation of the C4 amine, the latter inspired by the low cytotoxicity of the BlaS-related antimicrobial amicetin.^26^

## RESULTS AND DISCUSSION

### Improving the anti-malarial potency of blasticidin S through C6’ amidation

To determine whether C6’ amidation could be a productive route to improving the anti-malarial efficacy of BlaS, we screened a panel of eight amides (**Fig. 2A**) that have been investigated previously for their anti-microbial activity.^24^ These were assayed for growth inhibition at 2 µM using the drug-sensitive 3D7 parasite line in a 48-hour, fluorescence-based assay for nucleic acid content.^27^ Amidation products from ammonia (**2a**; generating the natural product P10),^28^ methylamine (**2b**), or ethylamine (**2c**) offered no improvement over the carboxylate of BlaS (**Fig. 2B**). Longer aliphatic amides (**2d-f**) modestly improved growth inhibition, whereas the presence of a hydrophilic hydroxyl group (**2g**) was deleterious. Remarkably, amidation with phenethylamine (**2h**) dramatically enhanced growth inhibition. This initial survey suggested that increasing non-polar surface area off the C6’ amide improved anti-malarial activity and identified the phenethyl amide analog as a major improvement over BlaS.

**Figure 2.**
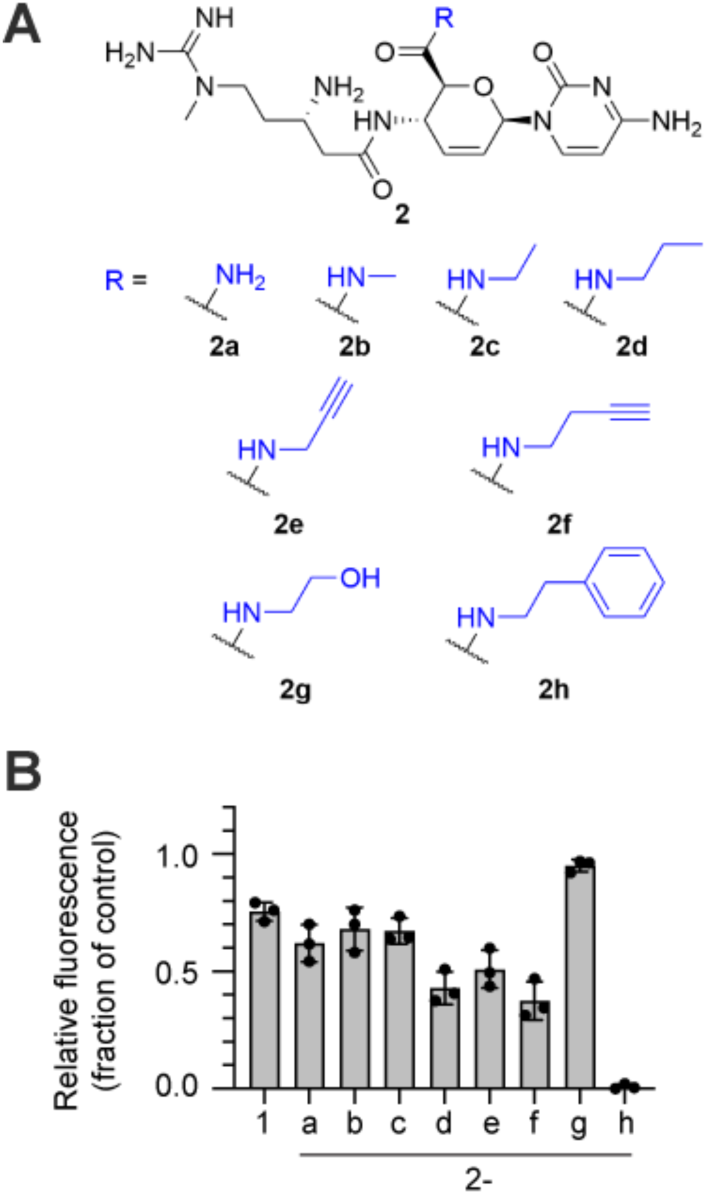
(**A**) Amide derivatives (**2**) of blasticidin S (**1**). (**B**) Single-concentration screen (2 µM) of parasite replication identifying phenethyl amide **2h** as an improved anti-parasitic compound. Bars represent mean values from three biological replicates with error bars indicating standard deviation. Values were normalized to cultures treated with vehicle (DMSO) and mefloquine

### Phenethyl SAR and linker optimization

We then conducted a structure-activity relationship (SAR) study around the C6’ phenethyl amide of **2h** to determine whether further improvements in potency are attainable. The synthetic approach mirrored that of our previous work (**Scheme 1**),^24^ activating the C6’ acid of **1** as the methyl ester (**3**) and protecting the nucleophilic beta-amine as the BOC carbamate (**4**). Diversification then proceeded by conversion of the protected methyl ester (**4**) to a series of corresponding new amides followed by deprotection (**5**) in a two-step process.

**Scheme 1.**
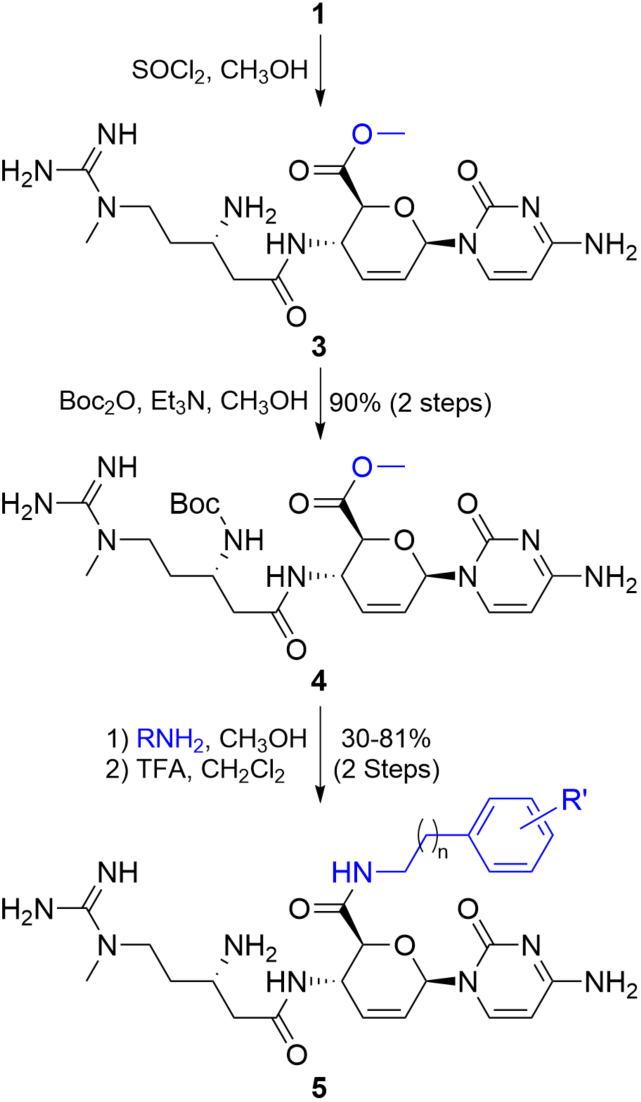
Synthetic approach to blasticidin S C6’ amide derivatives (**5**) diverging from protected ester intermediate **4**.

An initial slate of derivatives (**5a-c**) was synthesized to expand the SAR of **2h** by testing various electron donating and withdrawing groups (**Fig. 3A**). The 50% effective concentrations (EC_50_) for inhibition of parasite replication over one cycle (48 hours) were determined (**Fig. 3B, Table S1**), revealing 1.8- and 2.4-fold gains in potency for *para*-CF_3_ (**5b**) and -Cl (**5c**) substitutions, respectively. Reduced potency was observed for **5a**, which is consistent with the detrimental effects of introducing polarity that were observed for **2g**.

**Figure 3.**
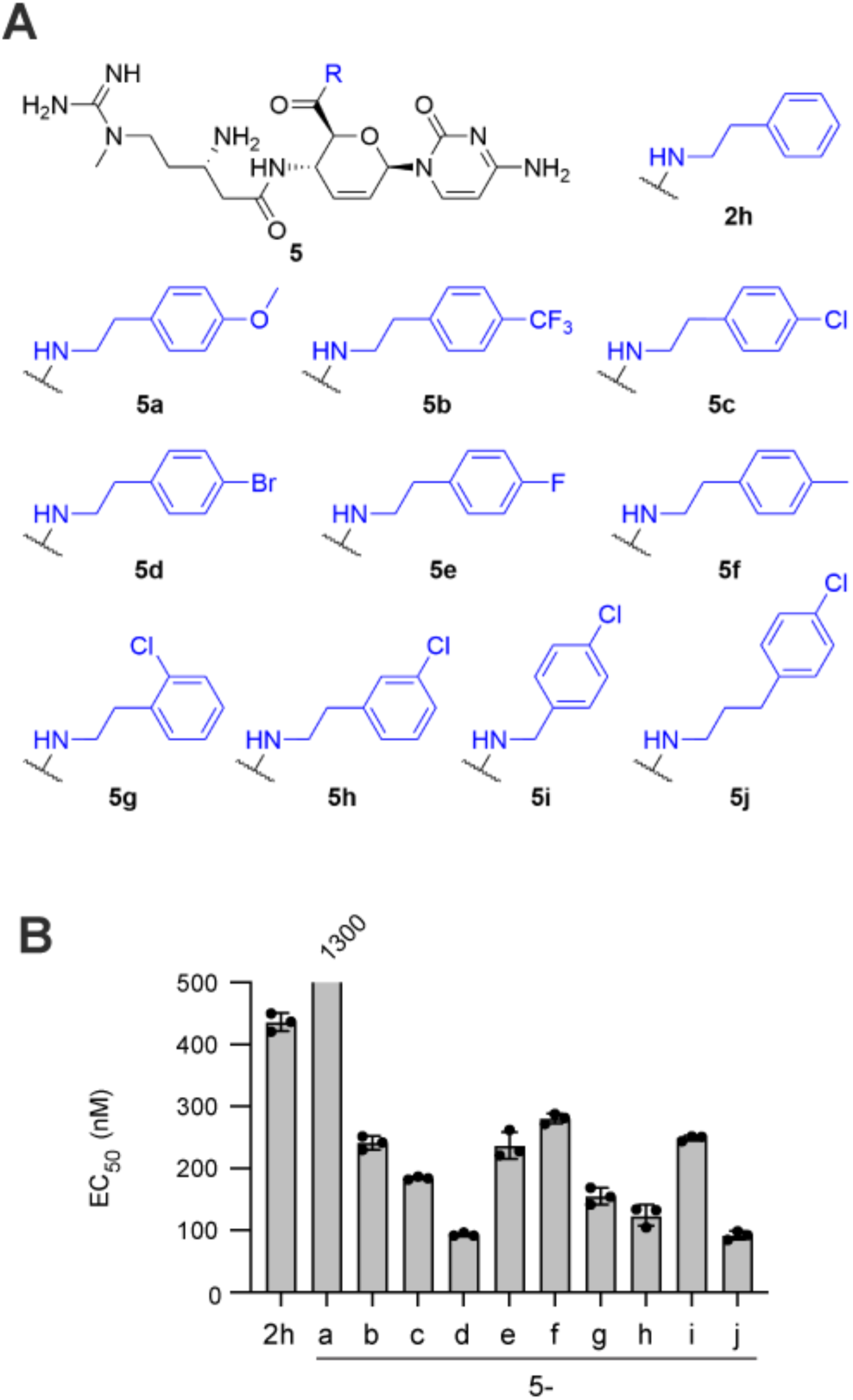
(**A**) Phenethyl amide derivatives **5a-j**. (**B**) EC50 values. Bars represent mean values from three biological replicates with error bars indicating standard deviation. The value for **5a** is off-scale and is indicated above the bar.

We next explored the SAR around **5c** by expanding the identity of the *para*-substitution (**5d-f**) and by shifting the position of chloride substitution to *ortho* (**5g**) and *meta* (**5h**). A two-fold improvement in EC_50_ was observed for the *p*-bromo substituent **5d**, with the other substitutions exhibiting either <2-fold decrease (**5h**,**g**) or a modest increase (**5e**,**f**) in EC_50_. Finally, we generated *p*-chloro analogs with one additional or one fewer methylene groups in the linker between the amide nitrogen and the aromatic ring. Increasing the linker length (**5j**) resulted in a two-fold enhancement of potency relative to **5c**, while reducing linker length (**5i**) had a negligible effect.

These studies yielded two compounds with sub-100 nM EC_50_ values against drug-sensitive *P. falciparum*: **5d** and **5j**. The EC_50_ value for BlaS for the 3D7 line is 6.6 µM (**Table 1**); thus, these analogs exhibit a ∼70-fold increase in potency over the parent compound. Compound **5j** was selected for further characterization as the chloride substituent was expected to have fewer metabolic liabilities than the brominated analog.

**Table 1:** Potencies of selected analogs and translation inhibitors against drug-sensitive (3D7) and multi-drug resistant (Dd2) parasite lines.

| Compound | Dd2 EC<sub>50</sub> (nM) | 3D7 EC<sub>50</sub> (nM) | EC<sub>50</sub> ratio 3D7:Dd2 |
| --- | --- | --- | --- |
| **5b** | 57 ± 2 | 240 ± 10 | 4.2 |
| **5c** | 40 ± 5 | 180 ± 10 | 4.5 |
| **5j** | 16 ± 1 | 92 ± 7 | 5.8 |
| **6a** | 71 ± 4 | 130 ± 10 | 1.8 |
| **6b** | 42 ± 6 | 87 ± 17 | 2.1 |
| **6c** | 150 ± 10 | 300 ± 60 | 2.0 |
| BlaS (**1**) | 7,100 ± 1,000 | 6,600 ± 1,200 | 0.9 |
| cycloheximide | 79 ± 4 | 97 ± 18 | 1.2 |
| anisomycin | 60 ± 5 | 67 ± 11 | 1.1 |

### Acylation at C4: blastimidines as a second optimization route

We have recently described a series of blastimidines (blasticidin-amicetin hybrids) based on **2h** that are acylated on the C4 (cytosine) amine (**Fig. 4A**)^23^. Because amicetin displays substantially lower cytotoxicity than BlaS,^26^ we speculated that C4 acylation could provide a route to improving therapeutic index. Thus, four *p*-aminobenzoic acid (PABA) C4 acylation derivatives of **2h** (blastimidines **6a-d**) were tested for anti-malarial activity. Acylation with PABA (**6a**) and *N*-Me PABA (**6b**) enhanced the EC_50_ value by 3.4 and 5.0-fold, respectively (**Fig. 4B**). These gains were reversed, however, with larger acylating groups (*N*-acyl-PABA **6c**, *N*-α-Me-seryl PABA **6d**). The most potent analog, **6b**, exhibited a comparable anti-malarial potency to that of **5j** (87 and 92 nM, respectively; **Table 1**). To further evaluate their potential as anti-malarial lead compounds, both analogs were further investigated for cytotoxicity, cross-resistance, and mode of action.

**Figure 4.**
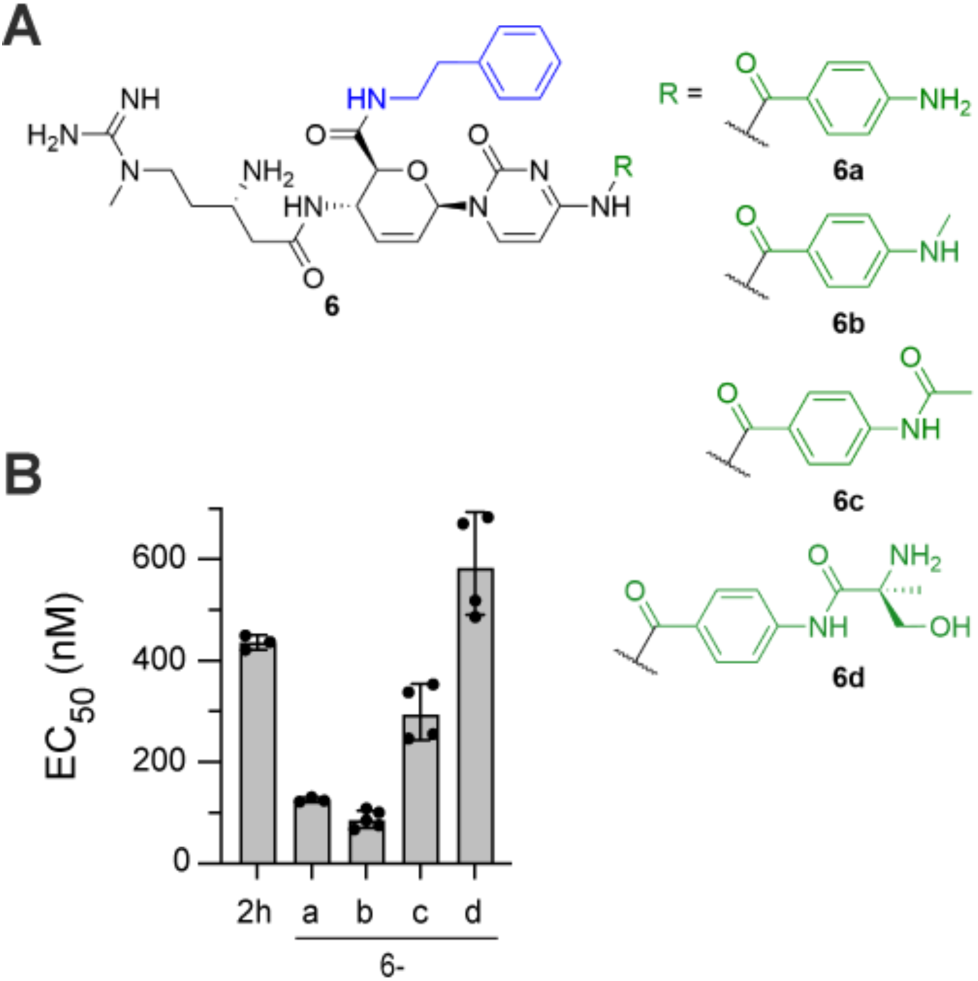
(**A**) Blastimidines **6a**–**6d** derived from C4 acylation of **2h**. (**B**) Mean EC50 values and standard deviation (error bars) from at least three biological replicates.

### Cytotoxicity

Blastimidine **6b** was previously shown to have negligible cytotoxicity against Vero (African green monkey kidney) cells following a 24 hour exposure (50% cell cytotoxicity (CC_50_) value > 256 µg/mL).^23^ To assess the cytotoxicity of **5j** and compare it to that of **6b**, both compounds were assayed with HepG2 cells, a human hepatocyte cell line that is frequently used to assess the cytotoxicity of anti-malarial leads. HepG2 cultures were exposed to compounds for 24 hours and cell viability was assayed using a tetrazolium reduction assay. In two independent experiments, the CC_50_ for both compounds was above 128 µg/mL, the highest concentration tested (corresponding to 142 and 128 µM for **5j** and **6b**, respectively), suggesting minimal mammalian toxicity in this assay. Combining this value with 50% effective concentrations (EC_50_) for inhibition of parasite replication collected above, we estimate the *P. falciparum:*HepG2 selectivity index to be >1,000 for both compounds. This effect is likely due to differences in interactions with mammalian and parasite targets.

### BlaS analogs exhibit enhanced potency against a multi-drug resistant parasite line

To assess the potential for cross-resistance with ubiquitous modes of drug resistance, we compared the potency of selected analogs against the multidrug-resistant Dd2 parasite line to that of the sensitive 3D7 line (**Table 1**). Interestingly, **5j** and **6b** were 6- and 2-fold more potent, respectively, against Dd2. Expanding the test set to include C6’ analogs **5b** and **5c** and blastimidines **6a** and **6c** revealed a class-specific effect, with the C6’ analogs exhibiting greater enhancements in potency than the blastimidines. Surprisingly, this effect was not observed with BlaS or the translation elongation inhibitors cycloheximide and anisomycin (**Table 1**), indicating that it is elicited by our modifications to the blasticidin S scaffold and is not a general property of translation inhibitors.

The Dd2 line has mutations in the *P. falciparum* chloroquine (CQ) resistance transporter (PfCRT) that render it less sensitive to chloroquine, a duplication in the gene that codes for *P. falciparum* multidrug resistance transporter 1 (PfMDR1) that confers mefloquine resistance, and a mutation in the dihydrofolate reductase-thymidylate synthase gene that provides resistance against anti-folates^29^. As transport processes are operative in the resistance mechanisms of chloroquine and mefloquine in the Dd2 line,^29^ it is possible that the modifications to BlaS C6’ and C4 groups have increased the affinity of these compounds for parasite transporters. PfCRT confers chloroquine resistance by transporting the drug from the food vacuole lumen to the cytosol.^30^ By analogy, enhanced efflux of **5j** from the food vacuole to the site of cytosolic ribosome inhibition could explain its enhanced potency. We employed the CQ resistance-reversing agent verapamil,^31^ which blocks PfCRT-mediated efflux from the food vacuole, to ask whether PfCRT contributes to the enhanced potency of **5j**. EC_50_ values for **5j** and CQ were determined in the presence and absence of 1 µM verapamil. While verapamil sensitized parasites to CQ as expected (EC_50_ ratio −/+ verapamil of 2.7 ± 0.4), there was essentially no effect with **5j** (EC_50_ ratio −/+ verapamil of 0.88 ± 0.06). Thus, PfCRT-mediated efflux does not seem to be responsible for this phenomenon. PfMDR1 amplification is also unlikely to be the cause; this food vacuole membrane transporter is thought to mediate resistance by transporting solutes across the food vacuole membrane into the lumen.^32^ Such a mechanism, if operative with **5j**, would be expected to reduce rather than enhance its potency against a cytosolic target.

### 5j and 6b inhibit parasite protein synthesis with improved kinetics over blasticidin S

We sought to assess whether **5j** and **6b** retain the mode of action of BlaS, namely inhibition of the cytosolic ribosome. Protein synthesis was quantified by incubating cultured parasites with methionine-free medium containing the clickable methionine analog homopropargylglycine (HPG).^33^ Cultures of trophozoites (24-30 hpi) were pre-treated with BlaS, **5j**, **6b** or the elongation inhibitors cycloheximide and anisomycin at 10x EC_50_ for 1, 2, 3 or 4 hours, at which point HPG was added for an additional two hours, yielding total incubation times of 3 to 6 hours. At the end of the labeling period, parasites were fixed and permeabilized, and HPG-labeled proteins were rendered fluorescent by click addition of an azide-conjugated fluorophore. Fluorescence was quantified by flow cytometry.

Treatment of parasites with cycloheximide and anisomycin resulted in a steady state of translation inhibition within one hour of exposure (**Fig. 5A**). Onset of translation inhibition by **5j** and **6b** occurred more slowly; however, the magnitude of inhibition was comparable to that of cycloheximide after three hours of pre-treatment. Surprisingly, little inhibition of translation was observed for BlaS even after a four hour pre-incubation period, which implies that either the rate of diffusion into the infected erythrocyte or the on-rate for ribosome binding is substantially slower than that for **5j** and **6b**. The data for **5j** and **6b** are consistent with translation inhibition as the primary mode of action and reveal that the rates of ribosome inhibition *in situ* are substantially improved over that of the parent compound.

**Figure 5.**
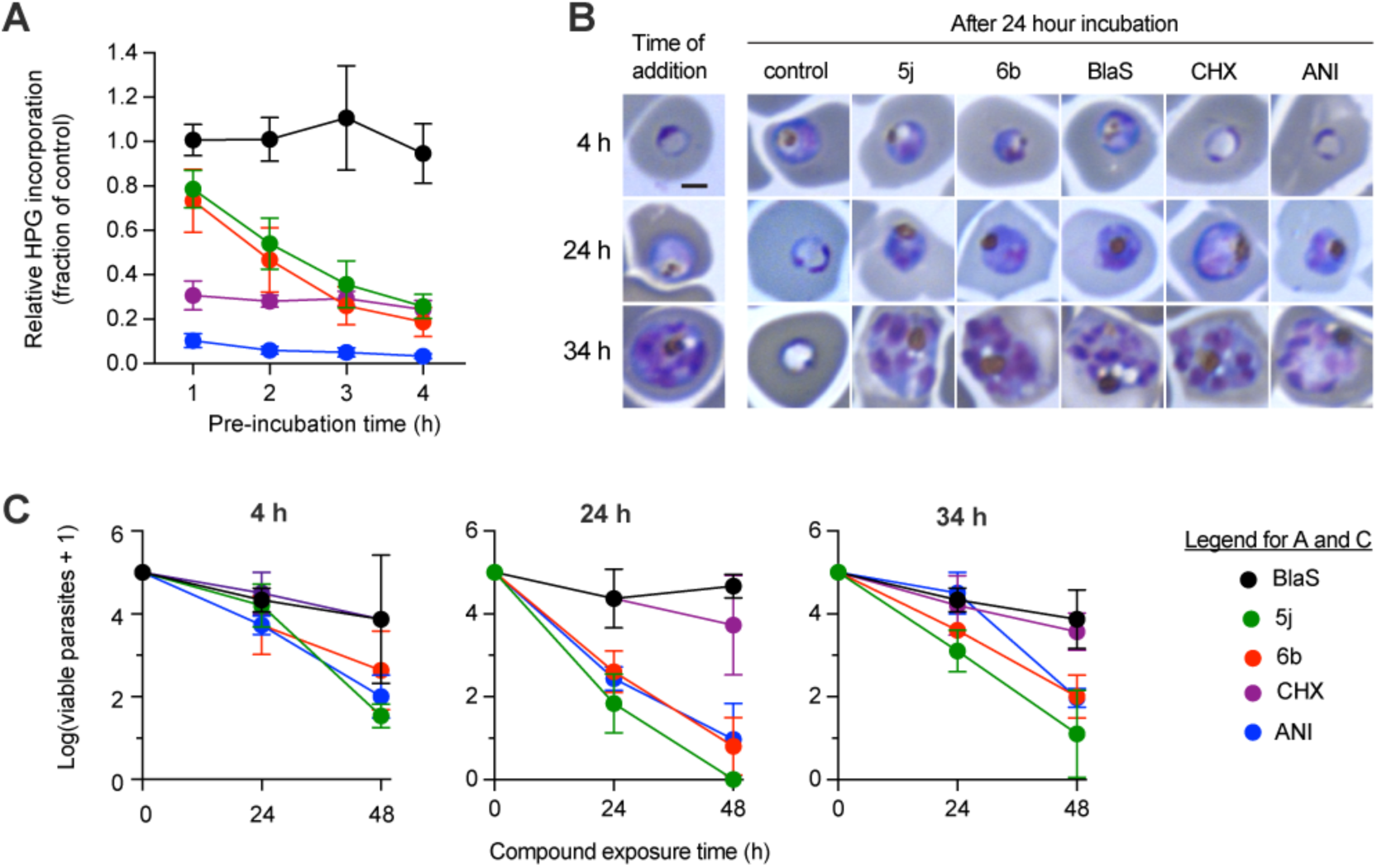
(**A**) Effect of ribosome inhibitors on homopropargylglycine (HPG) incorporation by cultured parasites. “Pre-incubation time” is the duration of exposure to compound prior to HPG addition, which was followed by an additional two hour incubation. Total incubation times therefore range from 3-6 h. Data are means and standard deviations from three independent experiments. (**B**) Stage specificity of action of ribosome inhibitors. Compounds were added at three timepoints (4, 24 and 34 hpi) and cultures were incubated for 24 hours. Representative images from Giemsa-stained smears are shown. Similar results were observed in three independent experiments. Bar, 2 µm. (**C**) Speed of parasite killing by ribosome inhibitors added at three developmental timepoints (4, 24 and 34 hpi). Cultures were incubated with compound for 24 or 48 hours followed by washout and serial dilution. Data are means and standard deviations from three independent experiments.

### Effect on asexual development and parasiticidal activity

We next evaluated the effects of BlaS, **5j**, **6b** and the translation elongation inhibitors cycloheximide and anisomycin on asexual parasite development. Highly synchronized ring-stage parasites (two hour invasion window) were generated and test compounds were added at 4, 24, and 34 hours post-invasion (hpi). These times were selected to assess the impact of compounds on the development of young ring, young trophozoite, and mid-schizont stages, respectively. In each case, parasite development was assessed 24 after compound addition by inspection of Giemsa-stained smears.

When added at 4 hpi, cycloheximide and anisomycin arrested parasites growth at the ring stage, an observation that is consistent with a prior report.^34^ In contrast, parasites treated with BlaS, **5j** and **6b** progressed to young trophozoites, which were distinguished from ring stages by the larger, rounded shape, darker Giemsa staining, and the presence of a hemozoin crystal (**Fig. 5B**). Inspection of cultures after an additional 24 hour incubation period (48 hours total exposure) revealed no further development of compound-treated cultures beyond that described above, whereas the control culture had completed a replication cycle and presented high ring parasitemia. When test compounds were added at 24 hpi, all appeared to arrest any further development and parasites remained at the young trophozoite stage, whereas the control culture had completed a replication cycle and consisted exclusively of ring-stage parasites. Likewise, when added at 34 hpi, all test compounds arrested schizont development (**Fig. 5B**).

These findings indicate that inhibitors of the cytosolic ribosomal translation can rapidly arrest parasite development from the early ring stage up to schizogony. The inability of BlaS, **5j**, and **6b** to arrest parasites at the ring stage (*i.e.*, when added at 4 hpi) may have its basis in the prior observation that BlaS, and likely **5j** and **6b** as well, relies on the plasmodial surface anion channel (PSAC) for entry into the infected erythrocyte^21^. PSAC is only expressed after ring-stage parasites transition to trophozoites, or at about 20 hours into the asexual replication cycle.^35^ Thus, the points at which BlaS, **5j** and **6b** arrest development align well with the onset of PSAC expression.

To determine the speed of killing by translation inhibitors, we treated parasites with test compounds at 10x EC_50_ at 4, 24, and 34 hours post-invasion (hpi) as described above. After 24 and 48 hours of treatment, compounds were washed out and 10^5^ parasites from the initial inoculum were serially diluted to determine the number of viable parasites. The results (**Fig. 5C**) reveal that compounds can be grouped into two clusters: 1) those that exhibited substantial cytotoxicity over 48 h at all times of addition (**5j**, **6b** and anisomycin) and 2) those that were predominantly cytostatic over the 48 h treatment period (BlaS, cycloheximide). As there is no obvious rationale for this trend on the basis of compound structure or site of ribosomal inhibition, its basis is presently unclear.

Parasites at 24 hpi were most sensitive to the effects of translation inhibitors. No viable parasites were obtained after 48 h treatment with **5j** (from two independent experiments), revealing a greater than 5-log reduction in viability, and **6b** and anisomycin elicited a 4-log reduction. The magnitude of reduction of viable parasitemia by these compounds is comparable to that found previously for chloroquine and mefloquine (between 4- and 5-log reduction after 48 hour treatment);^36^ thus, **5j** and **6b** exhibit relatively fast-acting parasiticidal activity when applied at 24 hpi. At 4 hpi, the reduction of viable parasitemia by **5j** and **6b** at 48 h was essentially identical to that after 24 h when added at 24 hpi, which is consistent with the approximately 20 h lag in efficacy observed in **Fig. 5B**. Parasiticidal activity was attenuated somewhat at 34 h, possibly due to reduced susceptibility of late-stage parasites; however, it is notable that **5j** remained highly parasiticidal, eliciting a 4-log reduction in parasitemia after 48 h treatment. Together, these studies provide a putative mechanism for uptake of **5j** and **6b** into infected erythrocytes *via* the PSAC channel and reveal dramatic improvements in parasiticidal activity over the parental compound BlaS.

### Modeling of 5j and 6b binding to the P*. falciparum* 80S ribosome

To gain insight into the structure–activity relationships observed across the C6′ amide and C4-acylated BlaS analogs, molecular docking was used to assess how specific synthetic modifications alter engagement with the *P. falciparum* ribosomal peptidyl transferase center (PTC). These modeling studies sought to rationalize how changes in aromatic surface area, linker length, and functional group identity translate into differential interaction networks within the PTC that are consistent with the measured EC₅₀ values. Molecular docking thus provides a structural lens through which the enhanced potency of the lead compounds **5j** and **6b** can be interpreted relative to the parent scaffold.

Compounds **6b** (**Fig. 6A**) and **5j** (**Fig. 6B**) exhibit interaction profiles that are consistent with their improved EC₅₀ values, with **6b** displaying the most favorable average predicted free energy of interaction among the compounds evaluated (**6b**, −8.6 kcal/mol; **5j**, −8.4 kcal/mol; BlaS, −7.9 kcal/mol). In contrast, docking of the parent compound BlaS (**Fig. 6C**) into the *P. falciparum* ribosomal PTC revealed a limited interaction network, consisting primarily of hydrogen bonding and a single π –π interaction. This comparatively sparse interaction profile is consistent with the weaker potency observed for BlaS relative to **5j** and **6b**. Importantly, comparison of the docked BlaS pose with the crystallographic BlaS orientation in the *Oryctolagus cuniculus* (rabbit) ribosome shows strong agreement in the positioning of the core ring system, with key interatomic distances ranging from ∼2.3–3.2 Å and a highly conserved overall orientation (**Fig. 6C**), supporting the validity of the docking protocol. However, divergence is observed in the orientation of the guanidinium group, which extends into a distinct cavity in the modeled *P. falciparum* structure. This region appears to provide greater spatial accommodation in the modeled ribosome than in the crystallographic reference, suggesting that the variation may arise from species-specific structural differences and/or conformational flexibility of the ribosomal environment that is not fully captured in static structures.

**Figure 6.**
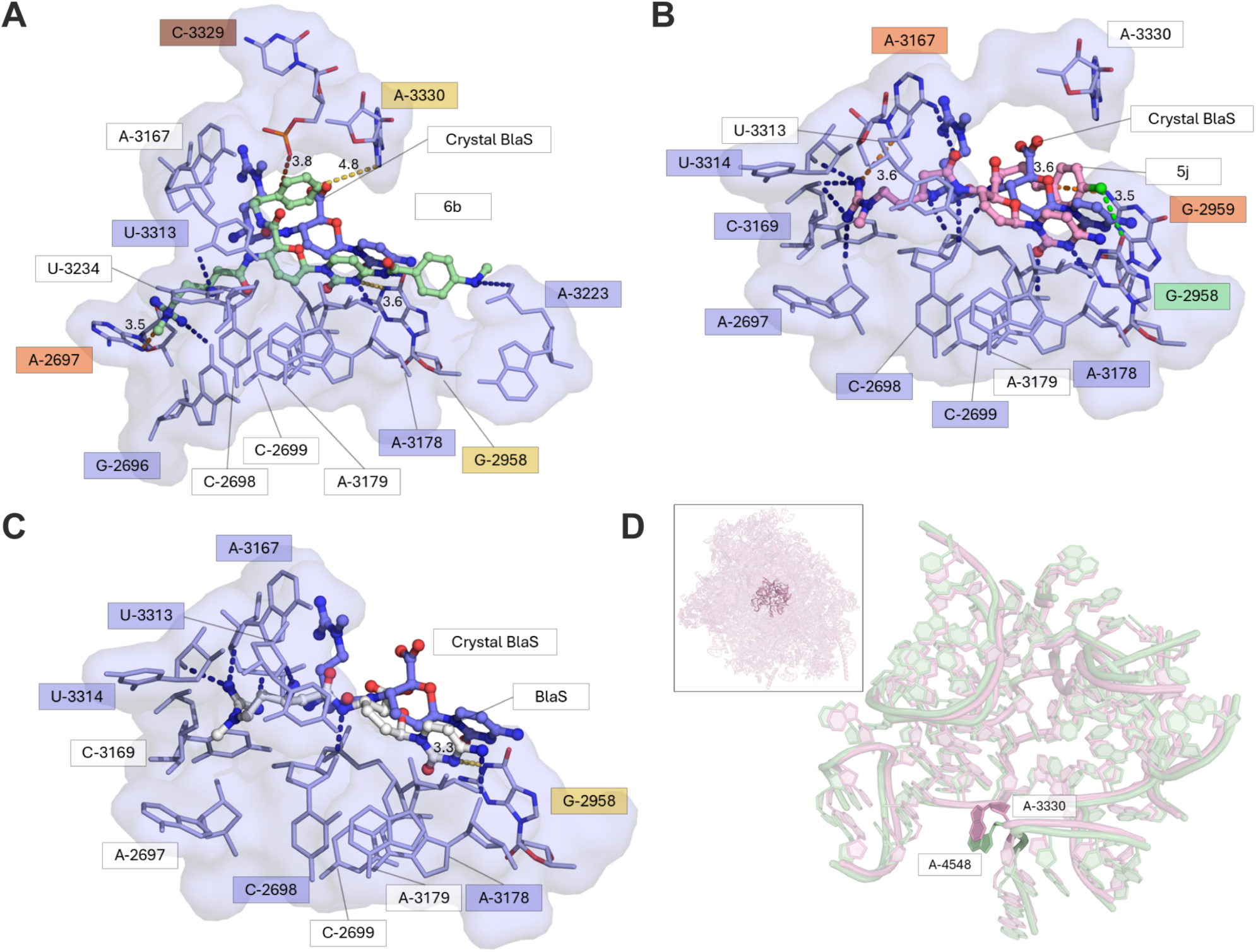
(**A-C**) Predicted binding poses of compounds **6b** (**A**; green), **5j** (**B**; pink), and **blasticidin S** (**C**; white) are shown docked into the *P. falciparum* 80S PTC (light blue; PDB ID: 5UMD), alongside the BlaS pose from the rabbit co-crystal structure (blue) for reference. Non-covalent interactions to nucleotides include π–π (pale yellow), cation–π (orange), halogen-mediated involving chlorine (green), hydrogen bonds (dark blue), and proximal (brown). Hydrogen bond distances are ≤4.0 Å and other bond distances are indicated in Å. (**D**) Alignment of the *P. falciparum* (light pink) and *O. cuniculus* (light green) PTCs highlighting nucleotides with species-specific conformational differences. Compared to A4548 in *O. cuniculus* (green), A3330 in *P. falciparum* (pink) exhibits a notably altered orientation and increased conformational flexibility, suggesting the potential for differential ligand engagement and species-selective binding. The inset shows the PTC (bold nucleotides) within the full 80S ribosome overlay.

The enhanced potency of **6b** is associated with the formation of an extensive network of noncovalent interactions within the A-site of the PTC of *P. falciparum*, including multiple cation–π and π–π interactions supplemented by hydrogen bonding (**Fig. 6A**). These contacts involve several key nucleotides lining the binding cavity (A2697, C3329, A3330, and G2958). The observed cation– and π-π interaction distances (3.5-3.6 Å) fall within established ranges for favorable noncovalent contacts,^37^ while a weaker π–π interaction (4.8 Å) and proximal interactions with C3329 are also present. Notably, these interactions are largely mediated by the two newly-introduced aromatic moieties in **6b**, highlighting the importance of expanded aromatic surface area in promoting productive engagement with the ribosomal A-site.

Compound **5j** also forms a richer interaction network than BlaS, combining hydrogen bonding and cation–π interactions with an additional halogen-mediated contact arising from the *para*-chloro substituent. The chlorine substituent is positioned to form a putative chlorine–oxygen interaction with nucleotide G2958, with interaction distance of 3.5 Å. (**Fig. 6B**). Cation–π interactions between **5j** and nucleotides A3167 and G2959 were also observed, with bond lengths of 3.6 Å. The halogen-enabled interaction introduces contact with a nucleotide that is also engaged by BlaS (**Fig. 6C**), and allows for additional cation-π interactions, providing a structural rationale for the enhanced potency of **5j** relative to the parent scaffold. While the conserved core ring structures of **5j** were positioned similarly to that of **6b**, the C6’ amide substituents differed in their orientations (**Fig. S1**) and formed distinctive interactions with the PTC.

Across the docked poses, A3330 emerged as a recurring interaction partner for both **5j** and **6b**, whereas BlaS contacted this nucleotide only sporadically: interaction with A3330 was observed in 1/9 BlaS poses versus 6/9 and 3/9 poses for **6b** and **5j**, respectively. Structural comparison revealed that A3330 adopts a markedly different orientation in the *P. falciparum* ribosome relative to the mammalian ribosome (**Fig. 6D**). This nucleotide (A4548 in *O. cuniculus* numbering) plays a critical role in peptide release, transpeptidation, and peptide bond formation,^38^ suggesting that preferential engagement of A3330 may contribute to both the potency and parasite selectivity observed for the optimized analogs.

## CONCLUSIONS

We have demonstrated that structural modifications at two sites of the BlaS scaffold can dramatically increase anti-malarial potency and selectivity. Compared to BlaS, the best-performing analogs **5j** and **6b** are ∼70-fold more potent against a drug-sensitive parasite line. Notably, potency is even greater against a multi-drug resistant line, yielding EC_50_ values as low as 16 nM. In cultured parasites, **5j** and **6b** inhibit protein translation substantially more rapidly than BlaS, a phenomenon that may be due to more effective transport through PSAC, a parasite-encoded, erythrocyte surface anion channel. Importantly, and in contrast to BlaS, **5j** and **6b** are fast-acting compounds with a 48-hour reduction in parasitemia of 10^4^- to10^5^-fold, values that are comparable to reductions observed with clinical anti-malarials. Molecular docking analysis reveals that **5j** and **6b** are likely to form more interactions with the *P. falciparum* ribosomal peptidyl transferase center. Together, our findings provide a roadmap for further optimization of potency and selectivity.

## MATERIALS AND METHODS

### Materials

Cycloheximide and anisomycin were obtained from Millipore Sigma. Met-free RPMI was generated in-house. Tris[(1-benzyl-1H-1,2,3triazol-4-yl)methyl]amine (TBTA) and CalFluor488 azide were obtained from Vector Laboratories. SYBR Green I and SYTOX AADvanced were obtained from ThermoFisher.

### P. falciparum culture

*Plasmodium falciparum* 3D7 and Dd2 were routinely cultured in RPMI in human O^+^ erythrocytes (Grifols Bio Supplies, Inc) at 2% hematocrit in RPMI 1640 medium supplemented with 0.37 mM hypoxanthine, 11 mM glucose, 27 mM sodium bicarbonate, 10 μg/mL gentamicin and 5 g/L Albumax I (Gibco). In addition, the medium was supplemented with 2 mM choline chloride, as this increases the multiplication rate by about 10%.^39^ Cultures were incubated at 37 °C in a 5% CO_2_ incubator with ambient O_2_ unless otherwise stated. Cultures were synchronized using the egress inhibitor ML10 and 5% (w/v) sorbitol as previously described.^40^

### Compound screening and EC_50_ assays

Growth inhibition assays were conducted in 200 µL volumes in 96 well plates. Parasite cultures were seeded at 3% parasitemia and 1% hematocrit and compounds were added from 500x DMSO stocks (BlaS analogs, cycloheximide, anisomycin) or from 200x water stocks (BlaS) in technical duplicate. Plates were incubated in a low-oxygen environment (5% O_2_, 5% CO_2_, 90% N_2_) for 48 hours and developed with the nucleic acid stain SYBR Green I as previously described.^27^ For initial screening, compounds were assessed at 2 µM. To determine EC_50_ values, a two-fold dilution series of 12 concentrations was assayed, with the EC_50_ values falling approximately in the midpoint of the concentration ranges. Fluorescence values (averages from technical duplicates) were fitted to a four-parameter sigmoidal dose-response curve by non-linear regression. Mean EC_50_ values and standard deviations were calculated from at least three independent experiments and are provided in **Table S1**.

### Homopropargylglycine protein synthesis assays

A culture of synchronized trophozoites at 6 – 10% parasitemia and 2% hematocrit was washed into Met-free RPMI, test compounds were added at a concentration of 10x EC_50_, and 200 µL aliquots were transferred to a 96-well plate. At hourly intervals up to four hours, HPG was added to 1 mM for an additional incubation period of two hours. At the end of the labeling period, cultures were washed twice with RPMI and fixed in 0.1% glutaraldehyde in phosphate-buffered saline (PBS). For the click reaction, fixed cells were permeabilized with 0.5% Triton X-100 for 20 minutes, washed twice with PBS, and incubated with 1 mM CuSO_4_, 0.1 mM TBTA, 2 mM sodium ascorbate and 100 µM CalFluor488 azide in 0.1 mg/mL bovine serum albumin/PBS as previously described.^41^ After incubation for 60 minutes at room temperature, cells were washed twice with PBS and taken up in 3 µM SYTOX AADvanced nuclear stain in PBS for 30 minutes. Cellular fluorescence was quantified on a Guava EasyCyte 5 flow cytometer. Parasite-infected erythrocytes were gated based on red SYTOX fluorescence and the median CalFluor 488 fluorescence for this population was determined using FlowJo 10.

### Development and speed of killing assays

To obtain synchronized parasites with a two-hour invasion window, egress-arrested parasites were generated by treating cultures with 100 nM ML10.^40^ Immediately following ML10 washout, cultures were adjusted to 4% hematocrit and rotated at 180 rpm on an orbital rotator with a 19 mm orbit diameter for one hour in a CO_2_ incubator at 37 °C. Remaining schizonts were removed by treatment with 5% sorbitol for 30 minutes at room temperature. The culture was adjusted to 3% parasitemia and 2% hematocrit and was rotated at 180 rpm throughout the experiment to promote synchronous development. Test compounds or DMSO vehicle (control) were added at 4, 24 and 34 hours post-invasion (hpi). Samples were collected at 24 h for preparation of Giemsa-stained smears. Images were collected on an AxioObserver equipped with a 100x/1.4NA objective and a color Axiocam MRc camera and contrast was adjusted using Adobe Photoshop.

Speed of killing assays were based on a previously described method for the determination of killing rates.^36^ Synchronized parasites were prepared as described in the above paragraph and test compounds or DMSO were added at 4, 24 and 34 hpi, with fresh medium (including test compounds) added after 24 h. After 24 and 48 hours of exposure, aliquots of culture corresponding to 10^5^ infected erythrocytes were washed to remove test compounds and were three-fold serially diluted in 96-well plates. Cultures were incubated for 21 days and parasite growth (or absence thereof) was confirmed by microscopy. Fold-change in viability was calculated based on the highest dilution containing parasites, compared with the DMSO control. Data are reported as means and standard deviations from three independent experiments.

### Cytotoxicity assays

HepG2-C3A cells were cultured in Dulbecco’s Modified Eagle’s Medium (DMEM) (Corning) containing 4.5 g/L glucose, 4mM L-glutamine, and sodium pyruvate and supplemented with 10% fetal bovine serum (FBS), 1% non-essential amino acids, 0.1% gentamicin sulfate, and 25 mM HEPES (herein referred to as DMEM-10). Cells were plated at 5,000 cells/well in a 96-well plate. Compound dilutions were prepared in DMEM-10. The vehicle (DMSO) was diluted to corresponding concentrations to serve as a control. Media was removed from plated HepG2-C3A cells and compound containing media was added. Cells were incubated at 37°C for 24 hours before CellTiter 96 AQueous One Solution Reagent was added to the cells according to the manufacturer’s protocol (Promega CellTiter 96 AQueous One Solution Cell Proliferation Assay). The cells were incubated for 2 hours at 37°C. Absorbance values were read at 490 nm using a Tecan Spark® Multimode Microplate Reader. Absorbance values were normalized to the corresponding DMSO control to determine impacts on cell viability.

### Molecular docking analysis

The 80S rabbit (*Oryctolagus cuniculus*) ribosomal unit with BlaS co-crystallized within the PTC (PDB ID: 7NWI)^18^ and *Plasmodium falciparum* 50S ribosomal subunits co-crystallized with mefloquine (PDB ID: 5UMD)^15^ and emetine (PDB ID: 3J79)^8^ were acquired. All ribosome structures had a resolution of 3.30 Å or less. PyMOL 3.1 and AutoDock Tools 1.5.7 were used to prepare the receptors for docking. The co-crystallized ligands were separated from the binding cavity, maintaining the relative atom coordinates. AutoDock Tools was used to add polar hydrogens to the receptors and compute gaisteiger charge. ChimeraX^42^ was used to prepare the ligands (BlaS, **6b**, **5j**, emetine and mefloquine) by adding polar hydrogens and removing nonpolar hydrogens. Water, small molecules, and tRNA were removed from the environment while polar hydrogens were added and the ions were retained. *P. falciparum* ribosome 50S and 30S subunits were overlaid and aligned to the 80S unit from *O. cuniculus* using PyMOL 3.1.6.1. The structures were processed to include only structural components within 40 Å of the center of the overlaid binding cavities to reduce computational load. Redocking was performed using GNINA 1.3. BlaS was redocked into the crystalized structure of *O. cuniculus* (PDB ID: 7NWI), emetine was redocked into *P. falciparum* (PDB ID: 6OKK) and mefloquine was redocked into *P. falciparum* (PDB ID: 5UMD) (**Fig. S2**). During redocking, the *O. cuniculus* ribosome co-crystallized with BlaS was used as a reference for comparing the success of A-site redocking of BlaS and derivative docks based on previous experiments with docking antibiotics into the bacterial PTC.^23, 24^ The redocking of BlaS was used to establish docking protocol coordinates and box size that could be used when docking into the aligned *P. falciparum* ribosome structures. The docking protocol box and coordinates used for all molecular docking in this work are center coordinates of 195.2, 192.6, 193.94 with a box size of 22 Å x 22 Å x 22 Å. The poses resulting from the docking of BlaS, **5j** and **6b** were separated and uploaded to Maestro to run fingerprint analysis using the Interaction Fingerprints feature. The fingerprint chart was exported using an inhouse python script. The nucleotides highlighted within the fingerprint chart were visualized in PyMOL 3.1.6.1. Representative docking poses shown in **Fig. 6** were selected by consensus across independent docking runs (PDB IDs: 5UMD and 3J79), retaining orientations consistent with the predominant pose clusters, conserved core placement relative to BlaS, and recurrent positioning of newly introduced substituents (*e.g*., aromatic rings and chlorine groups).

## ASSOCIATED CONTENT

### Supporting Information

Table S1 (EC_50_ values); Figures S1 & S2 (docking analysis); Synthetic methods; Figures S3 – S22 (^1^H and ^13^C NMR spectra) (PDF)

## AUTHOR INFORMATION

### Author Contributions

The manuscript was written through contributions of all authors. All authors have given approval to the final version of the manuscript. ‡These authors contributed equally.

### Notes

The authors declare no competing financial interest.

## ACKNOWLEDGMENTS

This work was supported by an Interdisciplinary Team-building Pilot Grant from the Center for Emerging, Zoonotic, and Arthropod-borne Pathogens at Virginia Tech. We are grateful to Nancy Vogelaar and the Virginia Tech Center for Drug Discovery Screening Lab for use of a multi-well plate liquid handling system.

## ABBREVIATIONS

ACT: artemisinin combination therapy
BlaS: blasticidin S
CQ: chloroquine
DMSO: dimethylsulfoxide;
HPG: homopropargylglycine
hpi: hours post-invasion
PABA: *para*-aminobenzoic acid
PfCRT: *P. falciparum* chloroquine resistance transporter
PfMDR1: *P. falciparum* multi-drug resistance transporter 1
PSAC: plasmodial surface anion channel
PTC: peptidyl transferase center.

