## Supporting Information for "Development of semisynthetic blasticidin S analogs with potent and fast-killing anti-malarial activity"

**Table S1:** $EC_{50}$ values from 48-hour *P. falciparum* growth inhibition assays.

| Compound | Phenyl subst.[1] | Linker[2] | $EC_{50}$ (nM) mean ± SD | $EC_{50}$ ratio[3] BlaS:X | $EC_{50}$ ratio[3] **2h**:X | $EC_{50}$ ratio[3] **5c**:X |
|---|---|---|---|---|---|---|
| BlaS | na[4] | na | 6,600 ± 1,200 | na | na | na |
| *C6' amides* | | | | | | |
| **2h** | none | 2 | 440 ± 20 | 15 | na | 0.4 |
| **5a** | *para*-OMe | 2 | 1,300 + 300 | 5 | 0.3 | 0.1 |
| **5b** | *para*-$CF_3$ | 2 | 240 ± 10 | 28 | 1.8 | 0.8 |
| **5c** | *para*-Cl | 2 | 180 ± 10 | 37 | 2.4 | na |
| **5d** | *para*-Br | 2 | 93 ± 3 | 71 | 4.7 | 1.9 |
| **5e** | *para*-F | 2 | 240 ± 20 | 28 | 1.8 | 0.8 |
| **5f** | *para*-Me | 2 | 280 ± 10 | 24 | 1.6 | 0.6 |
| **5g** | *orthro*-Cl | 2 | 120 ± 20 | 55 | 3.7 | 1.5 |
| **5h** | *meta*-Cl | 2 | 160 ± 10 | 41 | 2.8 | 1.1 |
| **5i** | *para*-Cl | 1 | 250 ± 10 | 26 | 1.8 | 0.7 |
| **5j** | *para*-Cl | 3 | 92 ± 7 | 72 | 4.8 | 2.0 |
| *Blastimidines* | | | | | | |
| **6a** | none | 2 | 130 ± 10 | 51 | 3.4 | na |
| **6b** | none | 2 | 87 ± 17 | 76 | 5.1 | na |
| **6c** | none | 2 | 300 ± 60 | 22 | 1.5 | na |
| **6d** | none | 2 | 590 ± 100 | 11 | 0.7 | na |

[1] Substituent on the C6' phenyl ring
[2] Number of methylene groups between the C6' amide and the phenyl ring
[3] Fold change in potency of compound X against the referents BlaS, **2h** or **5c**
[4] na, not applicable

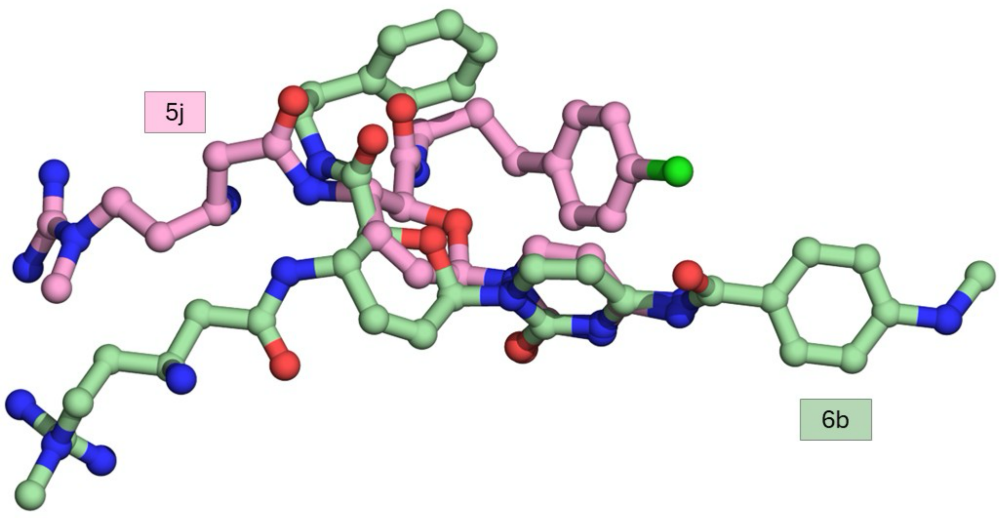


**Figure S1.** Comparison of docked poses of compounds **6b** and **5j** in the *P. falciparum* peptidyl transfer center. Representative docked poses of **6b** (pale green) and **5j** (pink) are overlaid to illustrate conserved core ring placement and differences due to C6' and C4 substituents.

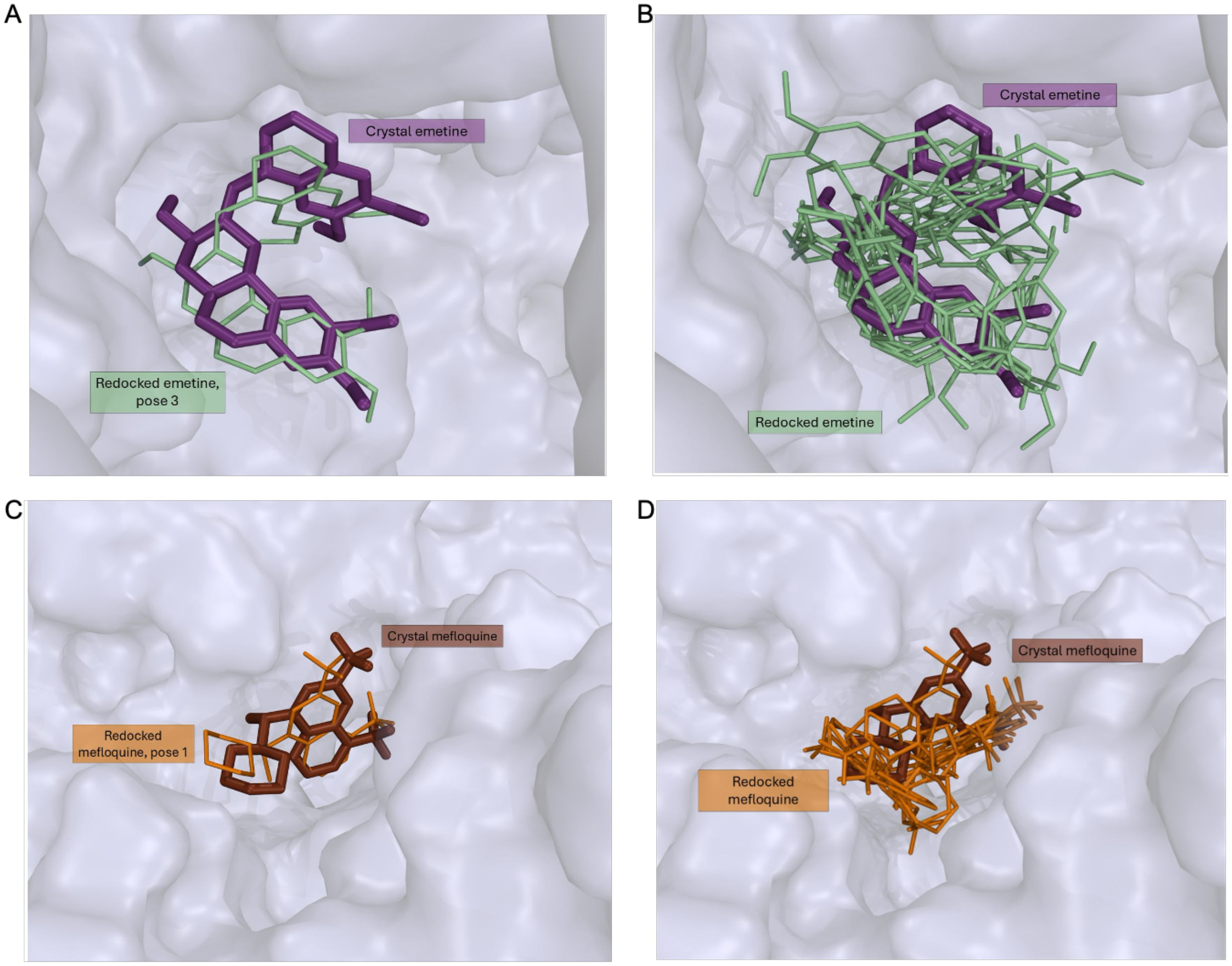


**Figure S2.** Docking protocol validation by redocking of reference ligands into the *P. falciparum* ribosome. Redocked poses of emetine (**A, B**; pale green) and mefloquine (**C, D**; orange) are shown overlaid with their crystallographic conformations (PDB ID: 3J79 - emetine, purple; PDB ID: 5UMD - mefloquine, brown). The best-ranked poses (Pose 3 for emetine (**A**) and Pose 1 for mefloquine (**C**)) reproduced the experimental binding modes with RMSD values of 1.6 and 1.8 Å, respectively.

**Synthetic methods**

Blasticidin S hydrochloride was purchased from Diagnocine. Unless otherwise specified, all reagents, solvents, and media components were purchased commercially and used as received from Sigma Aldrich, Fisher Scientific, or Oakwood Chemical. Deionized water was obtained from the house deionized water system. All synthetic reactions were stirred with a magnetic stir bar under a nitrogen atmosphere unless otherwise stated.

Specific rotations were obtained on a Jasco P-2000 polarimeter. $^1$H NMR spectra were recorded on a Bruker Avance II 500 MHz spectrometer, Agilent U4-DD2 400 MHz spectrometer, or Bruker Avance III 600 MHz spectrometer. Chemical shifts are reported in parts per million (ppm) using the solvent resonance as an internal standard ($CD_3OD$ 3.31 ppm, $D_2O$ 4.79 ppm). Data are reported as follows: chemical shift, multiplicity (s=singlet, d=doublet, t=triplet, q=quartet, m=multiplet), coupling constants (Hz), and number of protons. Proton decoupled $^{13}$C NMR were recorded on a Bruker Avance II 500 MHz ($^{13}$C 125 MHz) spectrometer or an Agilent U4-DD2 400 MHz ($^{13}$C 100 MHz) spectrometer. Chemical shifts are reported in ppm using the solvent resonance as an internal standard ($CD_3OD$ 49.0 ppm). NMR signals attributed to the counteranions (formate, trifluoroacetate) for salts are not tabulated but included in yield calculations. High resolution mass spectra were obtained on an Agilent Technologies 6220 TOF LC/MS, a Waters Synapt Q-TOF G2, or Thermo Exploris 120 HESI Orbitrap MS in the Department of Chemistry or the VT-Mass Spectrometry Incubator at the Virginia Polytechnic Institute and State University. Automated flash chromatography was performed using a Biotage Selekt system using water (unmodified, 0.1% (v/v) formic acid, or 0.1% (v/v) TFA) as solvent A and acetonitrile (unmodified, 0.1% (v/v) formic acid, or 0.1% (v/v) TFA) as solvent B. Commercial C18 cartridges were purchased from Biotage.

**Methyl (2*S*,3*S*,6*R*)-6-(4-amino-2-oxopyrimidin-1(2*H*)-yl)-3-((*S*)-3-((*tert*-butoxycarbonyl)amino)-5-(1-methylguanidino)pentanamido)-3,6-dihydro-2*H*-pyran-2-carboxylate mono formic acid salt (4).**

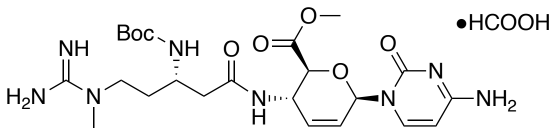


Blasticidin S hydrochloride **1** (502 mg, 1.09 mmol) was suspended in anhydrous MeOH (65.0 mL) and cooled in an ice bath with stirring. Thionyl chloride (2.40 mL, 33.1 mmol) was added slowly, and to the cold suspension with stirring. The resulting solution was warmed to rt, and after stirring for 18 h, was concentrated under a stream of dry nitrogen and then concentrated under vacuum. The resulting solids were dissolved in methanol (3 mL) and concentrated. This process was repeated and the solids further dried under vacuum to yield **3** as the trihydrochloride salt which was used directly in the next reaction.

Compound **3** trihydrochloride was dissolved in anhydrous methanol (12.2 mL) and cooled (ice bath). Triethylamine (0.35 mL, 2.5 mmol) was added to the stirring mixture, followed by Boc anhydride (0.33 mL, 1.4 mmol) and the mixture was warmed to rt. After stirring for 18 h, the mixture was concentrated to a white solid residue. This residue was dissolved in deionized water (3 mL) and purified using automated flash chromatography (C18, 30 g, 0-60% B, solvents modified with 0.1% formic acid) to yield **4** (572 mg, 90%, 2 steps) as a white amorphous solid. Spectral data were in accord with those previously reported.[1]

**(2*S*,3*S*,6*R*)-6-(4-amino-2-oxopyrimidin-1(2*H*)-yl)-3-((*S*)-3-amino-5-(1-methylguanidino)pentanamido)-*N*-phenethyl-3,6-dihydro-2*H*-pyran-2-carboxamide trihydrochloride (2h)**

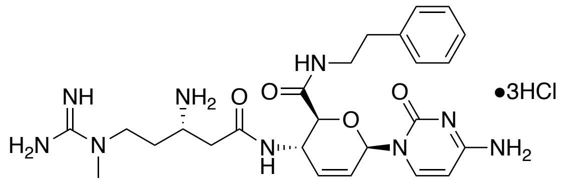


The mono-formic acid salt of **4** (77.4 mg, 0.133 mmol) was dissolved in a 1:6 (v/v) solution of anhydrous methanol to phenethylamine (1.75 mL) and stirred for 18 h in a sealed vial. The reaction was concentrated under a stream of nitrogen and then vacuum to yield a white solid, which was dissolved in a minimal amount of deionized water and purified using automated flash chromatography (C18, 30 g, 0-50% B, 0.1% formic acid) to yield the mono formic acid salt of **2h-BOC** as an amorphous white solid.

The mono formic acid salt of **2h-BOC** was dissolved in a stirring solution of 1:1 $CH_2Cl_2$/TFA (4.0 mL) while cooling (ice bath). The solution was warmed to rt over 2 h and then concentrated under vacuum. The resulting residue was treated with benzene and then the benzene removed under vacuum. This benzene treatment was repeated two more times before the residue was dissolved in a minimal amount of deionized water and purified using automated flash chromatography (C18, 30 g, 0-50% B, 0.1% formic acid) to yield the mono formic acid salt of **2h**, which was converted to the trihydrochloride by dissolution in 0.3% (v/v) conc. $HCl_{(aq.)}$ in methanol (5 mL) and concentration, this dissolution/concentration being repeated a total of three times. Drying under vacuum yielded **2h** as the trihydrochloride salt (50.8 mg, 75%, 2 steps). Spectral data were in accord with those previously reported.[1]

**(2*S*,3*S*,6*R*)-6-(4-Amino-2-oxopyrimidin-1(2*H*)-yl)-3-((*S*)-3-amino-5-(1-methylguanidino)pentanamido)-*N*-(4-methoxyphenethyl)-3,6-dihydro-2*H*-pyran-2-carboxamide mono formic acid di-TFA salt (5a)**

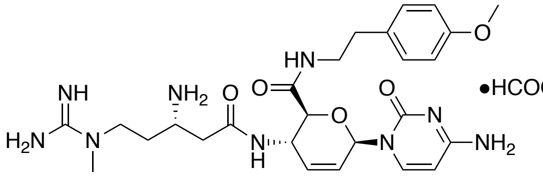


The mono-formic acid salt of **4** (80.4 mg, 0.138 mmol) was dissolved in a 1:4 (v/v) solution of anhydrous methanol to 4-methoxy phenethylamine (1.25 mL) and stirred for 18 h in a sealed vial. The mixture was diluted with ether (5 mL) and filtered to yield a white residue which was dissolved in a minimal amount of deionized water with 0.1% formic acid and purified using automated flash chromatography (C18, 30 g, 0-60% B, 0.1% formic acid) to yield the mono-formic acid salt of **5a-BOC** as an amorphous white solid.

The mono formic acid salt of **5a-BOC** was dissolved in a stirring solution of 1:1 $CH_2Cl_2$/TFA (2.0 mL) while cooling (ice bath). The solution was warmed to rt over 2 h and then concentrated under vacuum. The resulting residue was dissolved in a minimal amount of deionized water and purified using automated flash chromatography (C18, 30 g, 0-50% B, 0.1% formic acid) to yield **5a** (34.9 mg, 30%, 2 steps) as an amorphous white solid: $[\alpha]_D^{22}$ = +23° (*c* 1.7, $CH_3OH$); $^1H$ NMR (400 MHz, $CD_3OD$) δ 7.60 (d, *J* = 7.6 Hz, 1H), 7.13 – 7.06 (m, 2H), 6.84 – 6.78 (m, 2H), 6.59 (dt, *J* = 3.3, 1.8 Hz, 1H), 6.08 (dd, *J* = 10.1, 2.1 Hz, 1H), 6.04 – 5.97 (m, 1H), 5.90 – 5.83 (m, 1H), 4.84 – 4.80 (m, 1H), 4.19 (d, *J* = 9.4 Hz, 1H), 3.75 (m, 3H), 3.68 – 3.40 (m, 3H), 3.38 – 3.32 (m, 2H), 3.07 (s, 3H), 2.75 – 2.65 (m, 3H), 2.64 – 2.52 (m, 1H), 2.15 – 1.96 (m, 2H); $^{13}C$ NMR (100 MHz, $CD_3OD$) δ 171.3, 170.3, 166.3, 159.8, 158.4, 155.9, 144.1,

135.0, 132.2, 130.8, 127.2, 115.0, 96.9, 81.1, 77.5, 55.7, 48.4, 48.0, 46.8, 41.9, 38.2, 36.6, 35.4, 30.7; HRMS (ESI) calcd for $C_{26}H_{38}N_9O_5$ $[M+H]^+$ 556.2990, found 556.2991.

**(2*S*,3*S*,6*R*)-6-(4-Amino-2-oxopyrimidin-1(2*H*)-yl)-3-((*S*)-3-amino-5-(1-methylguanidino)pentanamido)-*N*-(4-(trifluoromethyl)phenethyl)-3,6-dihydro-2*H*-pyran-2-carboxamide mono formic acid di-TFA salt (5b).**

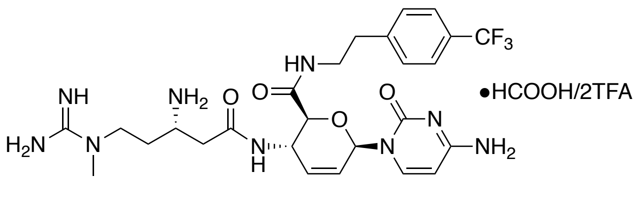


The mono-formic acid salt of **4** (88.8 mg, 0.152 mmol) was dissolved in a 1:4 (v/v) solution of anhydrous methanol to 4-trifluoromethyl phenethylamine (1.25 mL) and stirred for 18 h in a sealed vial. The mixture was diluted with ether (5 mL) and filtered to yield a white residue which was dissolved in a minimal amount of deionized water with 0.1% formic acid and purified using automated flash chromatography (C18, 30 g, 0-60% B, 0.1% formic acid) to yield the mono formic acid salt of **5b-BOC**.

The mono formic acid salt of **5b-BOC** was dissolved in a stirring solution of 1:1 $CH_2Cl_2$/TFA (3.0 mL) while cooling (ice bath). The solution was warmed to rt over 2 h and then concentrated under vacuum. The resulting residue was dissolved in a minimal amount of deionized water and purified using automated flash chromatography (C18, 30 g, 0-100% B, 0.1% formic acid) to yield **5b** (58.0 mg, 44%, 2 steps) as an amorphous white solid: $[\alpha]_D^{22}$ = +29° (*c* 2.9, $CH_3OH$); $^1H$ NMR (400 MHz, $CD_3OD$) δ 7.62 – 7.51 (m, 3H), 7.40 (d, *J* = 8.0 Hz, 2H), 6.59 (dt, *J* = 3.5, 1.8 Hz, 1H), 6.07 (dt, *J* = 10.2, 2.1 Hz, 1H), 5.98 (d, *J* = 7.5 Hz, 1H), 5.87 (ddd, *J* = 10.3, 2.7, 1.5 Hz, 1H), 4.85 – 4.80 (m, 1H), 4.19 (d, *J* = 9.4 Hz, 1H), 3.71 – 3.59 (m, 1H), 3.57 – 3.36 (m, 4H), 3.07 (s, 3H), 2.87 (t, *J* = 7.2 Hz, 2H), 2.71 (dd, *J* = 15.7, 4.5 Hz, 1H), 2.58 (dd, *J* = 15.7,

7.6 Hz, 1H), 2.06 (tq, $J$ = 10.7, 7.0 Hz, 2H); $^{13}$C NMR (100 MHz, $CD_3OD$) δ 171.4, 170.5, 166.7, 158.4, 156.6, 145.1, 143.9, 134.9, 130.6, 129.7 (q, $J$ = 32.1 Hz), 127.3, 126.3 (q, $J$ = 3.2 Hz) , 125.8 (q, $J$ = 271.4 Hz), 97.1, 81.2, 77.6, 48.4, 48.0, 46.8, 41.3, 38.3, 36.6, 36.0, 30.8; HRMS (ESI) calcd for $C_{26}H_{35}F_3N_9O_4$ $[M+H]^+$ 594.2759, found 594.2762.

**(2*S*,3*S*,6*R*)-6-(4-Amino-2-oxopyrimidin-1(2*H*)-yl)-3-((*S*)-3-amino-5-(1-methylguanidino)pentanamido)-*N*-(4-chlorophenethyl)-3,6-dihydro-2*H*-pyran-2-carboxamide mono formic acid di-TFA salt (5c).**

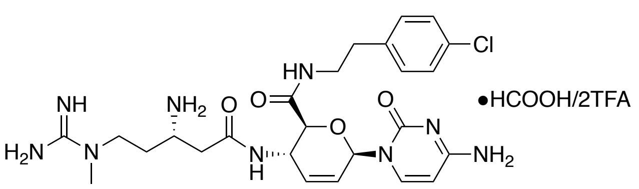


The mono-formic acid salt of **4** (81.0 mg, 0.125 mmol) was dissolved in a 1:4 (v/v) solution of anhydrous methanol to 4-chloro phenethylamine (1.25 mL) and stirred for 18 h in a sealed vial. The mixture was then diluted with ether (5 mL) and filtered to yield a white residue which was dissolved in a minimal amount of deionized water with 0.1% formic acid and purified using automated flash chromatography (C18, 30 g, 0-60% B, 0.1% formic acid) to yield the mono-formic acid salt of **5c-BOC**.

The mono formic acid salt of **5c-BOC** was dissolved in a stirring solution of 1:1 $CH_2Cl_2$/TFA (4.0 mL) while cooling (ice bath). The solution was warmed to rt over 2 h and then concentrated under vacuum. The residue was dissolved in a minimal amount of deionized water and purified using automated flash chromatography (C18, 30 g, 0-100% B, 0.1% formic acid) to yield **5c** (71.1 mg, 68%, 2 steps) as an amorphous white solid: $[\alpha]_D^{23}$ = +28° (*c* 3.6, $CH_3OH$); $^1$H NMR (400 MHz, $CD_3OD$) δ 7.60 (d, $J$ = 7.5 Hz, 1H), 7.30 – 7.21 (m, 2H), 7.21 – 7.15 (m, 2H),

6.62 – 6.56 (m, 1H), 6.08 (dt, $J$ = 10.3, 1.9 Hz, 1H), 6.02 (d, $J$ = 7.3 Hz, 1H), 5.87 (d, $J$ = 10.3 Hz, 1H), 4.85 – 4.80 (m, 1H), 4.21 (d, $J$ = 9.4 Hz, 1H), 3.67 – 3.58 (m, 1H), 3.56 – 3.45 (m, 2H), 3.43 – 3.33 (m, 2H), 3.07 (s, 3H), 2.77 (t, $J$ = 6.9 Hz, 2H), 2.70 (dd, $J$ = 15.6, 4.5 Hz, 1H), 2.60 (dd, $J$ = 15.7, 7.6 Hz, 1H), 2.12 – 1.99 (m, 2H); $^{13}C$ NMR (100 MHz, $CD_3OD$) δ 171.4, 170.4, 165.9, 158.3, 155.5, 144.3, 139.1, 135.1, 133.2, 131.5, 129.5, 127.0, 97.0, 81.1, 77.5, 48.5, 48.0, 46.7, 41.4, 38.3, 36.6, 35.5, 30.8; HRMS (ESI) calcd for $C_{25}H_{35}ClN_9O_4$ $[M+H]^+$ 560.2495, found 560.2496.

**(2*S*,3*S*,6*R*)-6-(4-Amino-2-oxopyrimidin-1(2*H*)-yl)-3-((*S*)-3-amino-5-(1-methylguanidino)pentanamido)-*N*-(4-bromophenethyl)-3,6-dihydro-2*H*-pyran-2-carboxamide tri-TFA salt (5d).**

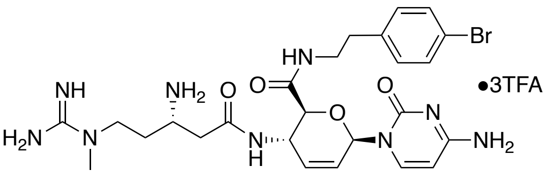


The mono-formic acid salt of **4** (76.7 mg, 0.132 mmol) was dissolved in a 1:5 (v/v) solution of anhydrous methanol to 4-bromo phenethylamine (1.5 mL) and stirred for 18 h in a sealed vial. The mixture was diluted with ether (5 mL) and filtered to yield a white residue which was dissolved in a minimal amount of deionized water with 0.1% formic acid and purified using automated flash chromatography (C18, 30 g, 0-100% B, 0.1% formic acid) to yield the mono-formic acid salt of **5d-BOC**.

The mono formic acid salt of **5d-BOC** was dissolved in a stirring solution of 1:1 $CH_2Cl_2$/TFA (3.5 mL) while cooling (ice bath). The solution was warmed to rt over 2 h and then concentrated under vacuum. The resulting residue was dissolved in a minimal amount of deionized water and purified using automated flash chromatography (C18, 30 g, 0-50% B, 0.1% TFA) to yield **5d** (100.7 mg, 81%, 2 steps) as an amorphous white solid: $[\alpha]_D^{23}$ = +20° (*c* 0.25, $CH_3OH$); $^1$H NMR (400 MHz, $CD_3OD$) δ 7.82 (d, *J* = 7.8 Hz, 1H), 7.46 – 7.38 (m, 2H), 7.18 – 7.10 (m, 2H), 6.59 (dt, *J* = 3.4, 1.9 Hz, 1H), 6.18 – 6.10 (m, 2H), 5.89 (ddd, *J* = 10.3, 2.7, 1.5 Hz, 1H), 4.86 – 4.82 (m, 1H), 4.21 (d, *J* = 9.4 Hz, 1H), 3.69 – 3.58 (m, 1H), 3.57 – 3.34 (m, 4H), 3.07 (s, 3H), 2.77 (t, *J* = 7.2 Hz, 2H), 2.71 (dd, *J* = 15.8, 4.5 Hz, 1H), 2.58 (dd, *J* = 15.8, 7.7 Hz, 1H), 2.14 – 1.97 (m, 2H); $^{13}$C NMR (100 MHz, $CD_3OD$) δ 171.4, 170.2, 161.8, 158.4, 148.9, 146.7, 139.6, 135.9, 132.5, 131.9, 126.0, 122.2, 96.0, 80.9, 77.5, 48.4, 48.0, 46.6, 41.4, 38.2, 36.6, 35.6, 30.7; HRMS (ESI) calcd for $C_{25}H_{35}BrN_9O_4$ $[M+H]^+$ 604.1990, found 604.2003.

**(2*S*,3*S*,6*R*)-6-(4-Amino-2-oxopyrimidin-1(2*H*)-yl)-3-((*S*)-3-amino-5-(1-methylguanidino)pentanamido)-*N*-(4-fluorophenethyl)-3,6-dihydro-2*H*-pyran-2-carboxamide tri-TFA salt (5e).**

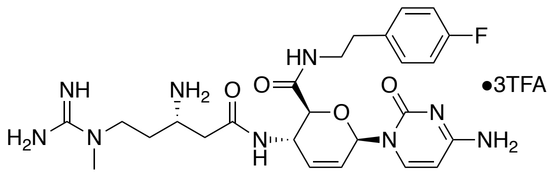


The mono-formic acid salt of **4** (79.1 mg, 0.136 mmol) was dissolved in a 1:5 (v/v) solution of anhydrous methanol to 4-fluoro phenethylamine (1.5 mL) and stirred for 18 h in a sealed vial. The mixture was then diluted with ether (5 mL) and filtered to yield a white residue which was dissolved in a minimal amount of deionized water with 0.1% formic acid and purified using automated flash chromatography (C18, 30 g, 0-100% B, 0.1% formic acid) to yield the mono-formic acid salt of **5e-BOC**.

The mono formic acid salt of **5e-BOC** was dissolved in a stirring solution of 1:1 $CH_2Cl_2$/TFA (2.6 mL) while cooling (ice bath). The solution was warmed to rt over 2 h and then concentrated under vacuum. The resulting residue was dissolved in a minimal amount of deionized water and purified using automated flash chromatography (C18, 30 g, 0-50% B, 0.1% TFA) to yield **5e** (82.9 mg, 69%, 2 steps) as an amorphous white solid: $[\alpha]_D^{24}$ = +18° (*c* 0.25, $CH_3OH$); $^1H$ NMR (400 MHz, $CD_3OD$) δ 7.83 (d, *J* = 7.8 Hz, 1H), 7.24 – 7.18 (m, 2H), 7.04 – 6.96 (m, 2H), 6.59 (dt, *J* = 3.4, 1.8 Hz, 1H), 6.17 – 6.12 (m, 2H), 5.89 (ddd, *J* = 10.3, 2.7, 1.6 Hz, 1H), 4.85 – 4.81 (m, 1H), 4.21 (d, *J* = 9.4 Hz, 1H), 3.68 – 3.59 (m, 1H), 3.58 – 3.42 (m, 2H), 3.42 – 3.33 (m, 2H), 3.07 (s, 3H), 2.78 (t, *J* = 7.3 Hz, 2H), 2.71 (dd, *J* = 15.8, 4.5 Hz, 1H), 2.58 (dd, *J* = 15.8, 7.7 Hz, 1H), 2.12 – 1.98 (m, 2H); $^{13}C$ NMR (100 MHz, $CD_3OD$) δ 170.0, 168.8, 161.6 (d, *J* = 243.0 Hz), 160.3, 156.9, 147.4, 145.3, 134.9 (d, *J* = 3.2 Hz), 134.5, 130.1 (d, *J* = 8.0 Hz), 124.6, 114.7 (d, *J* = 21.4 Hz), 94.5, 79.5, 76.0, 47.0, 46.6, 45.2, 40.3, 36.8, 35.2, 34.0, 29.3; HRMS (ESI) calcd for $C_{25}H_{35}FN_9O_4$ $[M+H]^+$ 544.2791, found 544.2807.

**(2*S*,3*S*,6*R*)-6-(4-Amino-2-oxopyrimidin-1(2*H*)-yl)-3-((*S*)-3-amino-5-(1-methylguanidino)pentanamido)-*N*-(4-methylphenethyl)-3,6-dihydro-2*H*-pyran-2-carboxamide tri-TFA salt (5f).**

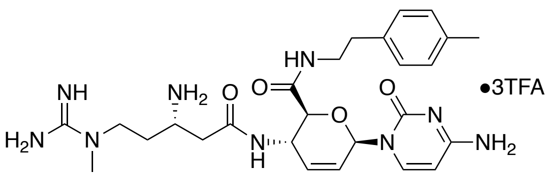


The mono-formic acid salt of **4** (78.0 mg, 0.134 mmol) was dissolved in a 1:5 (v/v) solution of anhydrous methanol to 4-methyl phenethylamine (1.5 mL) and stirred for 18 h in a sealed vial. The mixture was then diluted with ether (5 mL) and filtered to yield a white residue which was dissolved in a minimal amount of deionized water with 0.1% formic acid and purified using automated flash chromatography (C18, 30 g, 0-100% B, 0.1% formic acid) to yield the mono-formic acid salt of **5f-BOC**.

The mono formic acid salt of **5f-BOC** was dissolved in a stirring solution of 1:1 $CH_2Cl_2$/TFA (3.0 mL) while cooling (ice bath). The solution was warmed to rt over 2 h and then concentrated under vacuum. The resulting residue was dissolved in a minimal amount of deionized water and purified using automated flash chromatography (C18, 30 g, 0-50% B, 0.1% TFA) to yield **5f** (95.1 mg, 81%, 2 steps) as an amorphous white solid: $[\alpha]_D^{24}$ = +17° (*c* 0.25, $CH_3OH$); $^1H$ NMR (400 MHz, $CD_3OD$) δ 7.82 (d, *J* = 7.8 Hz, 1H), 7.08 (apparent s, 4H), 6.59 (dt, *J* = 3.4, 1.8 Hz, 1H), 6.18 – 6.10 (m, 2H), 5.89 (ddd, *J* = 10.3, 2.7, 1.5 Hz, 1H), 4.86 – 4.82 (m, 1H), 4.22 (d, *J* = 9.4 Hz, 1H), 3.66 – 3.59 (m, 1H), 3.59 – 3.42 (m, 2H), 3.40 – 3.34 (m, 2H), 3.07 (s, 3H), 2.77 – 2.68 (m, 3H), 2.58 (dd, *J* = 15.8, 7.7 Hz, 1H), 2.28 (s, 3H), 2.13 – 1.97 (m, 2H); $^{13}C$ NMR (100 MHz, $CD_3OD$) δ 171.4, 170.1, 161.8, 158.3, 149.0, 146.7, 137.1, 137.0, 135.9, 130.1, 129.7, 126.0, 96.0, 80.9, 77.4, 48.4, 48.0, 46.6, 41.9, 38.2, 36.6, 35.9, 30.7, 21.1. HRMS (ESI) calcd for $C_{26}H_{38}N_9O_4$ $[M+H]^+$ 540.3041, found 540.3045.

**(2*S*,3*S*,6*R*)-6-(4-Amino-2-oxopyrimidin-1(2*H*)-yl)-3-((*S*)-3-amino-5-(1-methylguanidino)pentanamido)-*N*-(2-chlorophenethyl)-3,6-dihydro-2*H*-pyran-2-carboxamide tri-TFA salt/0.75 mol 2-chlorophenethylamine adduct (5g).**

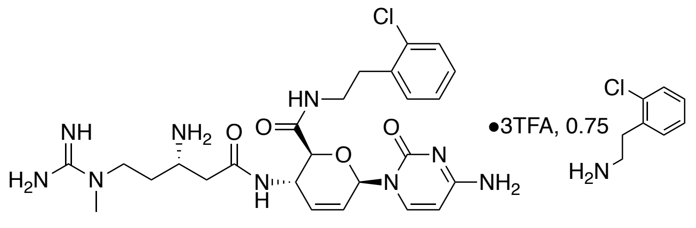

The mono-formic acid salt of **4** (75.0 mg, 0.129 mmol) was dissolved in a 1:5 (v/v) solution of anhydrous methanol to 2-chloro phenethylamine (1.5 mL) and stirred for 18 h in a sealed vial. The mixture was diluted with ether (5 mL) and filtered to yield a white residue which was dissolved in a minimal amount of deionized water with 0.1% formic acid and purified using automated flash chromatography (C18, 30 g, 0-100% B, 0.1% formic acid) to yield the mono-formic acid salt of **5g-BOC**.

The mono formic acid salt of **5g-BOC** was dissolved in a stirring solution of 1:1 $CH_2Cl_2$/TFA (2.5 mL) while cooling (ice bath). The solution was warmed to rt over 2 h and then concentrated under vacuum. The resulting residue was dissolved in a minimal amount of deionized water and purified using automated flash chromatography (C18, 30 g, 0-50% B, 0.1% TFA) to yield and inseparable mixture of **5g** plus 0.75 equiv. 2-chlorophenethylamine (56.7 mg, 50.2 mg **5g** as calculated by $^1$H NMR, 43%, 2 steps) as an amorphous white solid: $[\alpha]_D^{21}$ = +18° (*c* 0.25, $CH_3OH$); $^1$H NMR (400 MHz, $CD_3OD$) δ 7.84 (d, *J* = 7.8 Hz, 1H), 7.47 – 7.15 (m, 7H), 6.59 (dt, *J* = 3.4, 1.8 Hz, 1H), 6.19 – 6.10 (m, 2H), 5.89 (ddd, *J* = 10.3, 2.7, 1.5 Hz, 1H), 4.88 – 4.83 (m, 1H), 4.22 (d, *J* = 9.4 Hz, 1H), 3.68 – 3.58 (m, 1H), 3.58 – 3.39 (m, 4H), 3.23 – 3.09 (m, 3.5H), 3.07 (s, 3H), 2.95 (m, 1.5H), 2.71 (dd, *J* = 15.7, 4.5 Hz, 1H), 2.59 (dd, *J* = 15.7, 7.6 Hz, 1H), 2.13 – 1.98 (m, 2H); $^{13}$C NMR (125 MHz, $CD_3OD$) δ 170.0, 168.9, 161.0, 157.0, 148.5, 145.0, 136.4, 134.4, 134.2, 133.7, 131.0, 130.8, 129.5, 129.1, 128.8, 127.9, 127.3, 126.8, 124.7 (2C), 94.7, 79.5, 76.0, 47.0, 46.6, 45.3, 38.8, 38.6, 36.8, 35.2, 32.6, 31.1, 29.3; HRMS (ESI) calcd for $C_{25}H_{35}ClN_9O_4$ $[M+H]^+$ 560.2495, found 560.2493.

**(2*S*,3*S*,6*R*)-6-(4-Amino-2-oxopyrimidin-1(2*H*)-yl)-3-((*S*)-3-amino-5-(1-methylguanidino)pentanamido)-*N*-(3-chlorophenethyl)-3,6-dihydro-2*H*-pyran-2-carboxamide tri-TFA salt (5h).**

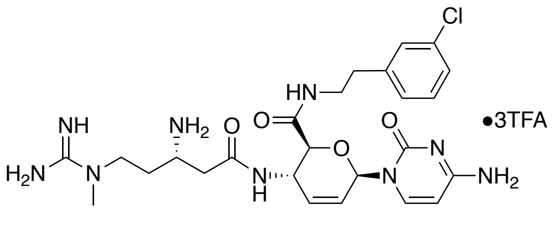


The mono-formic acid salt of **4** (78.8 mg, 0.129 mmol) was dissolved in a stirring solution of 1:5 (v/v) anhydrous methanol to 3-chloro phenethylamine (1.5 mL) and stirred for 18 h in a sealed vial. The mixture was diluted with ether (5 mL) and filtered to yield a white residue which was dissolved in a minimal amount of deionized water with 0.1% formic acid and purified using automated flash chromatography (C18, 30 g, 0-100% B, 0.1% formic acid) to yield the mono-formic acid salt of **5h-BOC**.

The mono formic acid salt of **5h-BOC** was dissolved in a stirring solution of 1:1 $CH_2Cl_2$/TFA (2.0 mL) while cooling (ice bath). The solution was warmed to rt over 2 h and then concentrated under vacuum. The resulting residue was dissolved in a minimal amount of deionized water and purified using automated flash chromatography (C18, 30 g, 0-50% B, 0.1% TFA) to yield **5h** (54.1 mg, 46%, 2 steps) as an amorphous white solid: $[\alpha]_D^{21}$ = +26° (*c* 0.25, $CH_3OH$); $^1H$ NMR (400 MHz, $CD_3OD$) δ 7.81 (d, *J* = 7.8 Hz, 1H), 7.33 – 7.23 (m, 2H), 7.20 (ddd, *J* = 7.8, 2.2, 1.3 Hz, 1H), 7.15 (dt, *J* = 7.2, 1.4 Hz, 1H), 6.59 (dt, *J* = 3.4, 1.8 Hz, 1H), 6.18 – 6.10 (m, 2H), 5.89 (ddd, *J* = 10.3, 2.7, 1.5 Hz, 1H), 4.86 – 4.80 (m, 1H), 4.22 (d, *J* = 9.4 Hz, 1H), 3.67 – 3.58 (m, 1H), 3.57 – 3.34 (m, 4H), 3.08 (s, 3H), 2.79 (t, *J* = 7.3 Hz, 2H), 2.72 (dd, *J* = 15.8, 4.5 Hz, 1H), 2.58 (dd, *J* = 15.8, 7.7 Hz, 1H), 2.14 – 1.96 (m, 2H); $^{13}C$ NMR (100 MHz, $CD_3OD$) δ 171.4, 170.2,

161.8, 158.4, 149.0, 146.6, 142.8, 135.8, 135.2, 131.1, 129.9, 128.4, 127.5, 126.1, 96.0, 80.9, 77.5, 48.4, 48.0, 46.7, 41.4, 38.2, 36.6, 35.9, 30.7; HRMS (ESI) calcd for $C_{25}H_{35}ClN_9O_4$ $[M+H]^+$ 560.2495, found 560.2508.

**(2*S*,3*S*,6*R*)-6-(4-Amino-2-oxopyrimidin-1(2*H*)-yl)-3-((*S*)-3-amino-5-(1-methylguanidino)pentanamido)-*N*-(4-chlorobenzyl)-3,6-dihydro-2*H*-pyran-2-carboxamide tri-TFA salt (5i).**

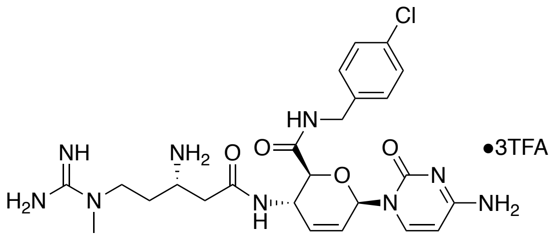


The mono-formic acid salt of **4** (76.7 mg, 0.126 mmol) was dissolved in a 1:5 (v/v) solution of anhydrous methanol to 4-chloro benzylamine (1.5 mL) and stirred for 18 h in a sealed vial. The reaction was then diluted with ether (5 mL) and filtered to yield a white residue which was dissolved in a minimal amount of deionized water with 0.1% formic acid and purified using automated flash chromatography (C18, 30 g, 0-100% B, 0.1% formic acid) to yield the mono-formic acid salt of **5i-BOC**.

The mono formic acid salt of **5i-BOC** was dissolved in a stirring solution of 1:1 $CH_2Cl_2$/TFA (2.0 mL) while cooling (ice bath). The solution was warmed to rt over 2 h and then concentrated under vacuum. The resulting residue was dissolved in a minimal amount of deionized water and purified using automated flash chromatography (C18, 30 g, 0-50% B, 0.1% TFA) to yield **5i** (42.9 mg, 38%, 2 steps) as an amorphous white solid: $[\alpha]_D^{21}$ = +23° (*c* 0.25, $CH_3OH$); $^1H$ NMR (400 MHz, $CD_3OD$) δ 7.84 (d, *J* = 7.8 Hz, 1H), 7.33 – 7.29 (m, 2H), 7.29 – 7.25 (m, 2H), 6.62 (dt, *J* = 3.4, 1.9 Hz, 1H), 6.15 (dt, *J* = 10.2, 2.0 Hz, 1H), 6.11 (d, *J* = 7.8 Hz, 1H), 5.90 (ddd,

$J$ = 10.3, 2.7, 1.5 Hz, 1H), 4.94 (dq, $J$ = 9.5, 2.9 Hz, 1H), 4.42 (d, $J$ = 15.2 Hz, 1H), 4.31 (d, $J$ = 9.2 Hz, 1H), 4.30 (d, $J$ = 15.8 Hz, 1H), 3.65 – 3.56 (m, 1H), 3.55 – 3.39 (m, 2H), 3.04 (s, 3H), 2.71 (dd, $J$ = 15.8, 4.5 Hz, 1H), 2.58 (dd, $J$ = 15.8, 7.5 Hz, 1H), 2.11 – 1.94 (m, 2H); $^{13}$C NMR (100 MHz, $CD_3OD$) δ 171.4, 170.4, 161.9, 158.3, 149.2, 146.7, 138.5, 135.9, 133.9, 130.1, 129.5, 126.1, 96.0, 80.9, 77.5, 48.4, 47.9, 46.7, 43.0, 38.1, 36.5, 30.7; HRMS (ESI) calcd for $C_{24}H_{33}ClN_9O_4$ $[M+H]^+$ 546.2339, found 546.2344.

**(2*S*,3*S*,6*R*)-6-(4-Amino-2-oxopyrimidin-1(2*H*)-yl)-3-((*S*)-3-amino-5-(1-methylguanidino)pentanamido)-*N*-(3-(4-chlorophenyl)propyl)-3,6-dihydro-2*H*-pyran-2-carboxamide tri-TFA salt (5j).**

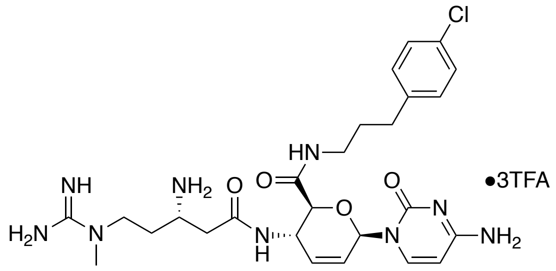


The mono-formic acid salt of **4** (75.8 mg, 0.124 mmol) was dissolved in a 1:5 (v/v) solution of anhydrous methanol to 4-chloro phenylpropylamine (1.5 mL) and stirred for 18 h in a sealed vial. The mixture was then diluted with ether (5 mL) and filtered to yield a white residue which was dissolved in a minimal amount of deionized water with 0.1% formic acid and purified using automated flash chromatography (C18, 30 g, 0-100% B, 0.1% formic acid) to yield the mono-formic acid salt of **5j-BOC**.

The mono formic acid salt of **5j-BOC** was dissolved in a stirring solution of 1:1 $CH_2Cl_2$/TFA (2.0 mL) while cooling (ice bath). The solution was warmed to rt over 2 h and then concentrated under vacuum. The resulting residue was dissolved in a minimal amount of deionized water and purified using automated flash chromatography (C18, 30 g, 0-60% B, 0.1% TFA) to yield **5j** (55.7 mg, 49%, 2 steps) as an amorphous white solid: $[\alpha]_D^{20}$ = +22° (*c* 0.3, $CH_3OH$); $^1H$ NMR (400 MHz, $CD_3OD$) δ 7.85 (d, *J* = 7.8 Hz, 1H), 7.28 – 7.22 (m, 2H), 7.21 – 7.15 (m, 2H), 6.60 (dt, *J* = 3.4, 1.8 Hz, 1H), 6.18 – 6.10 (m, 2H), 5.89 (ddd, *J* = 10.3, 2.7, 1.5 Hz, 1H), 4.89 – 4.87 (m, 1H), 4.22 (d, *J* = 9.4 Hz, 1H), 3.65 – 3.56 (m, 1H), 3.55 – 3.39 (m, 2H), 3.19 (qd, *J* = 13.5, 6.9 Hz, 2H), 3.06 (s, 3H), 2.70 (dd, *J* = 15.8, 4.5 Hz, 1H), 2.61 (t, *J* = 7.5 Hz, 2H), 2.57 (dd, *J* = 15.7, 7.7 Hz, 1H), 2.08 – 1.98 (m, 2H), 1.88 – 1.75 (m, 2H); $^{13}C$ NMR (100 MHz, $CD_3OD$) δ 171.3, 170.2, 161.9, 158.3, 149.2, 146.7, 141.8, 135.9, 132.6, 131.1, 129.4, 126.1, 95.9, 80.9, 77.6, 49.0, 48.0, 46.7, 39.8, 38.1, 36.6, 33.4, 31.8, 30.7; HRMS (ESI) calcd for $C_{26}H_{37}ClN_9O_4$ $[M+H]^+$ 574.2652, found 574.2650.

### Compound Spectra

**Figure S3.** $^{1}$H NMR (400 MHz, $CD_3OD$) Spectrum of **5a**.

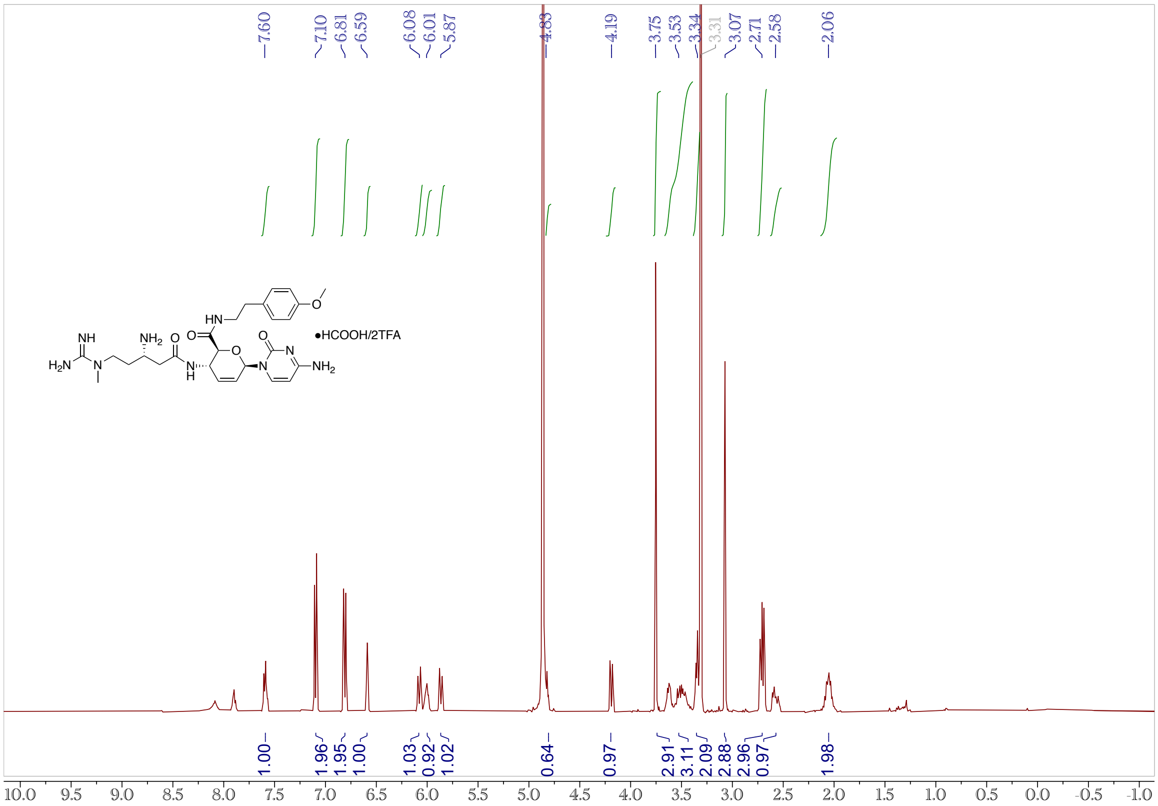


**Figure S4.** $^{13}$C NMR (100 MHz, $CD_3OD$) Spectrum of **5a**.

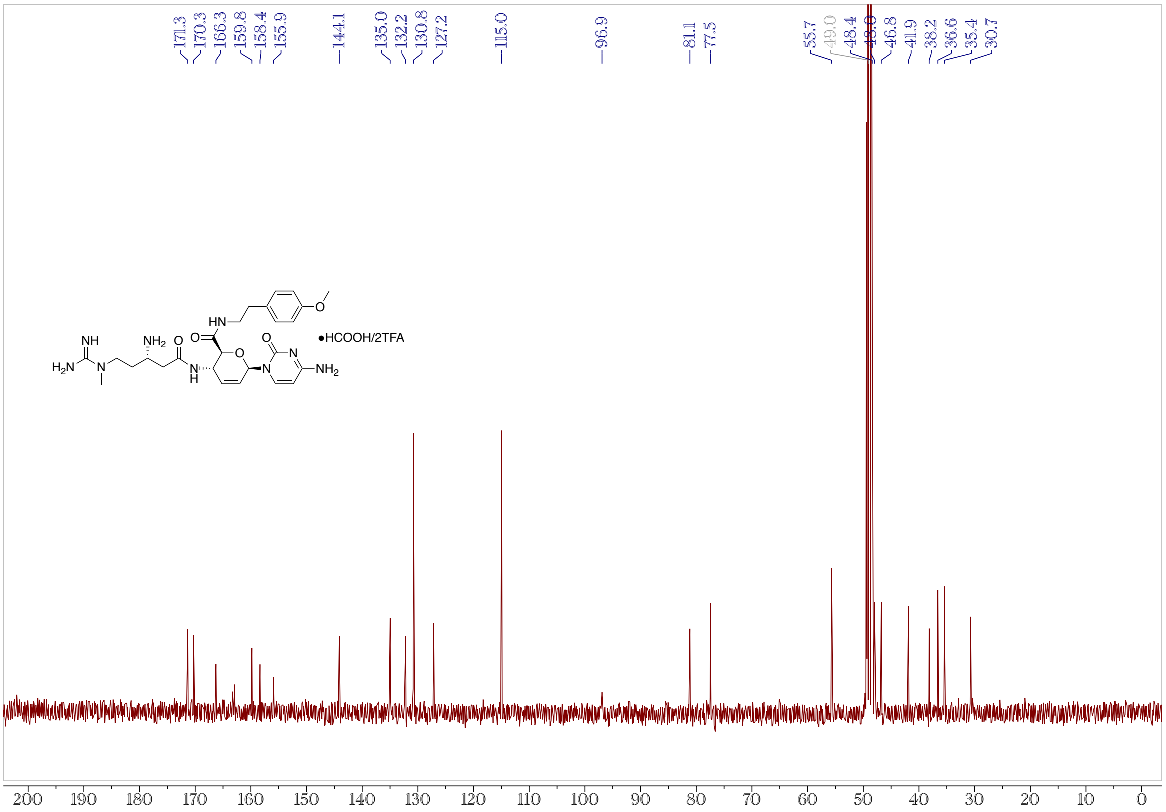

**Figure S5.** $^{1}H$ NMR (400 MHz, $CD_3OD$) Spectrum of **5b**.

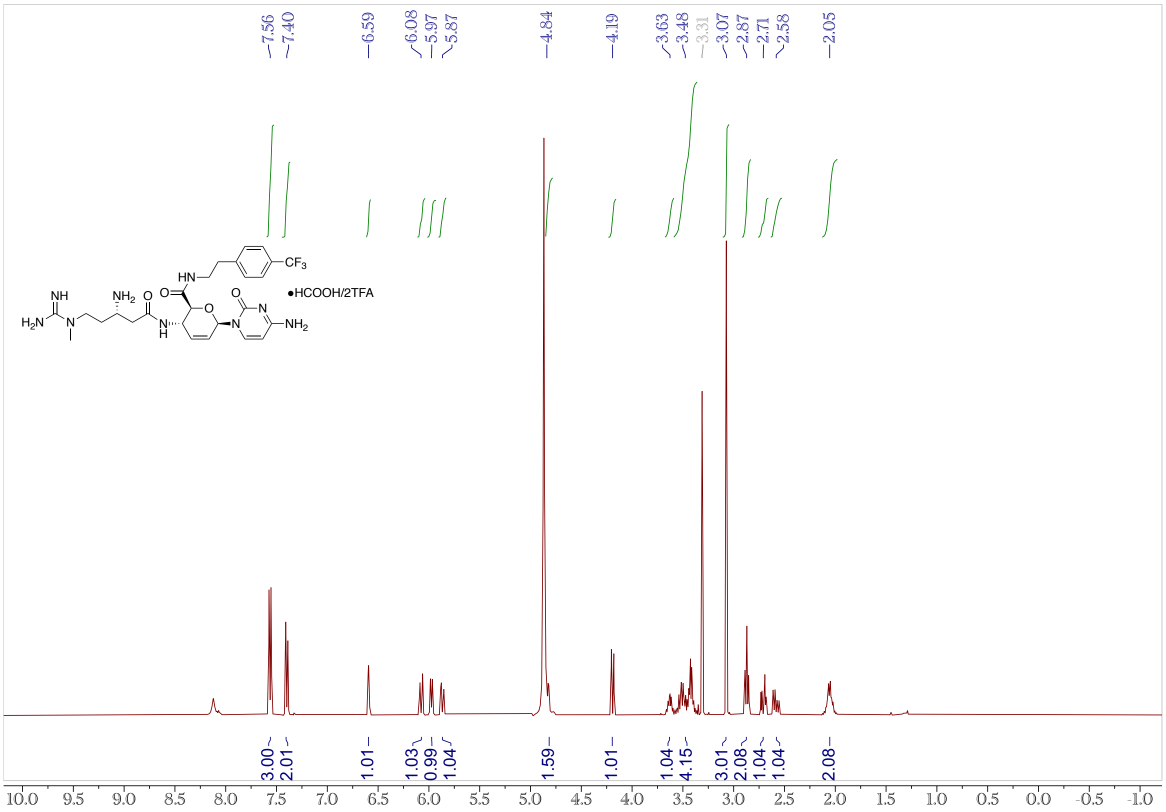


**Figure S6.** $^{13}C$ NMR (100 MHz, $CD_3OD$) Spectrum of **5b**.

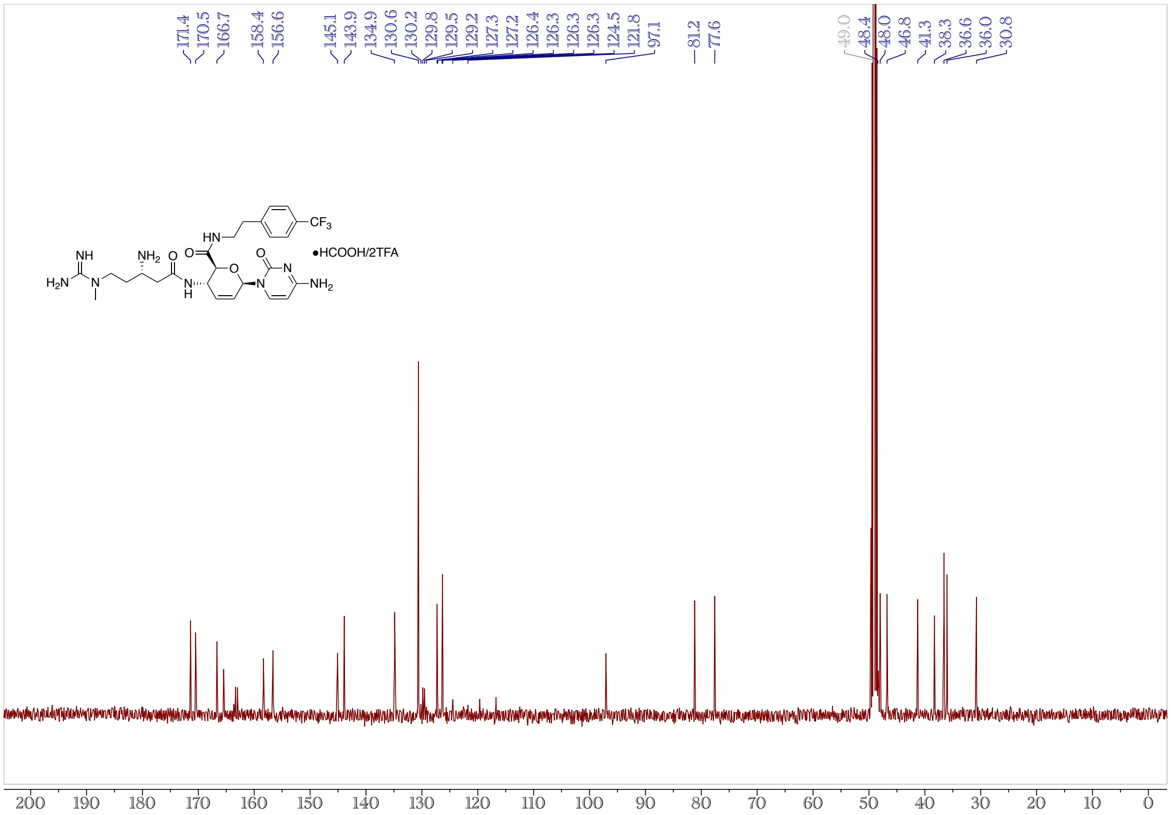

**Figure S7.** $^{1}$H NMR (400 MHz, $CD_3OD$) Spectrum of **5c**.

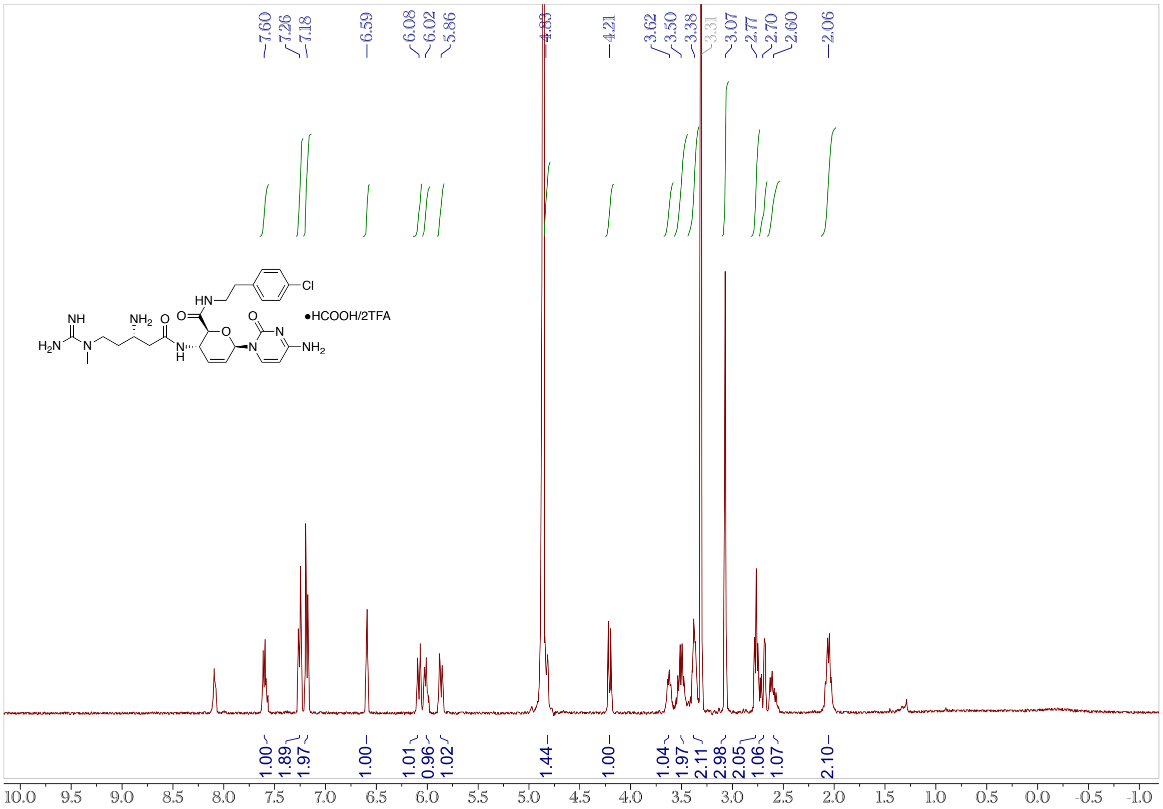

**Figure S8.** $^{13}$C NMR (100 MHz, $CD_3OD$) Spectrum of **5c**.

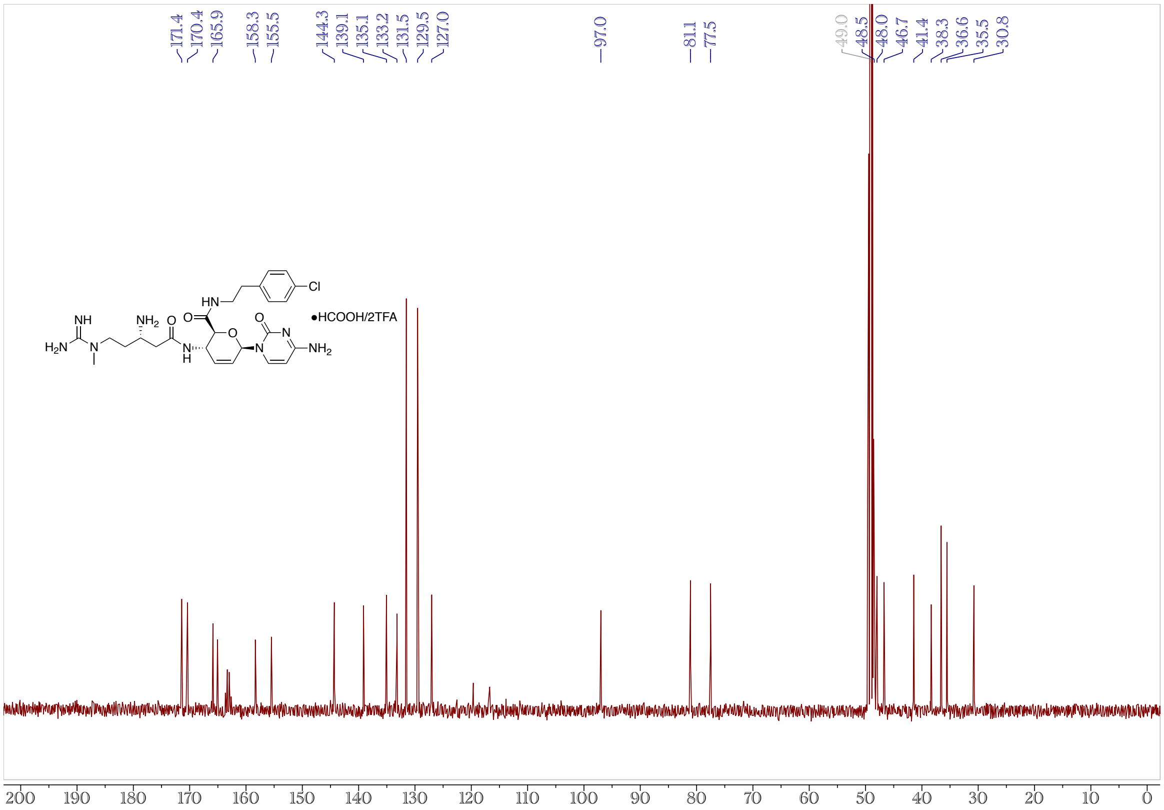

**Figure S9.** $^{1}$H NMR (400 MHz, $CD_3OD$) Spectrum of **5d**.

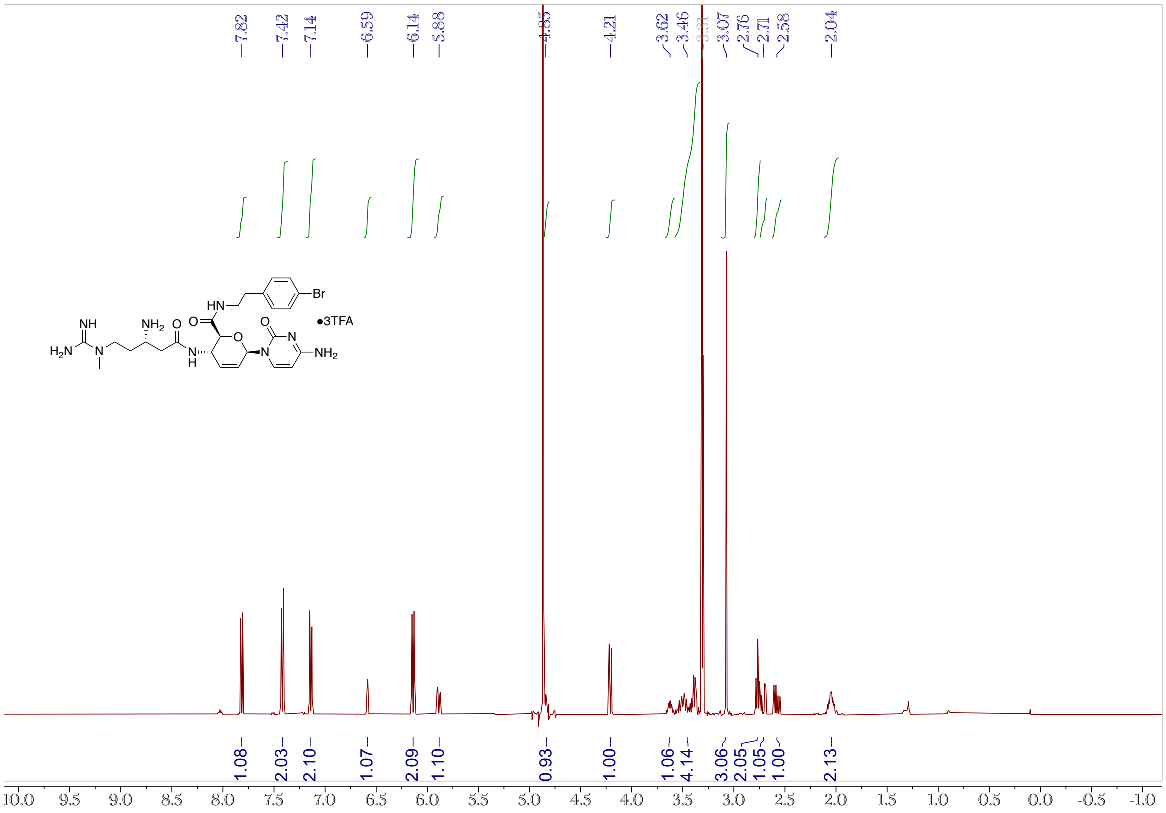


**Figure S10.** $^{13}$C NMR (100 MHz, $CD_3OD$) Spectrum of **5d**.

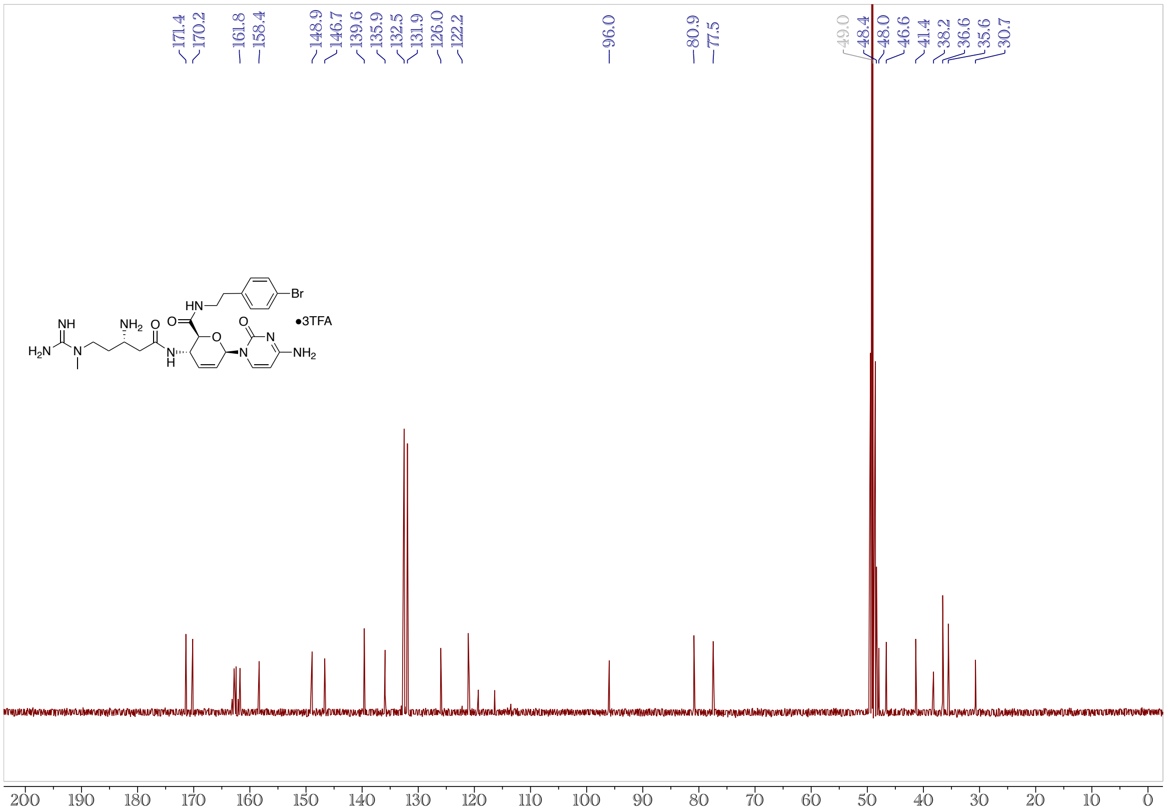

**Figure S11.** $^1$H NMR (400 MHz, $CD_3OD$) Spectrum of **5e**.

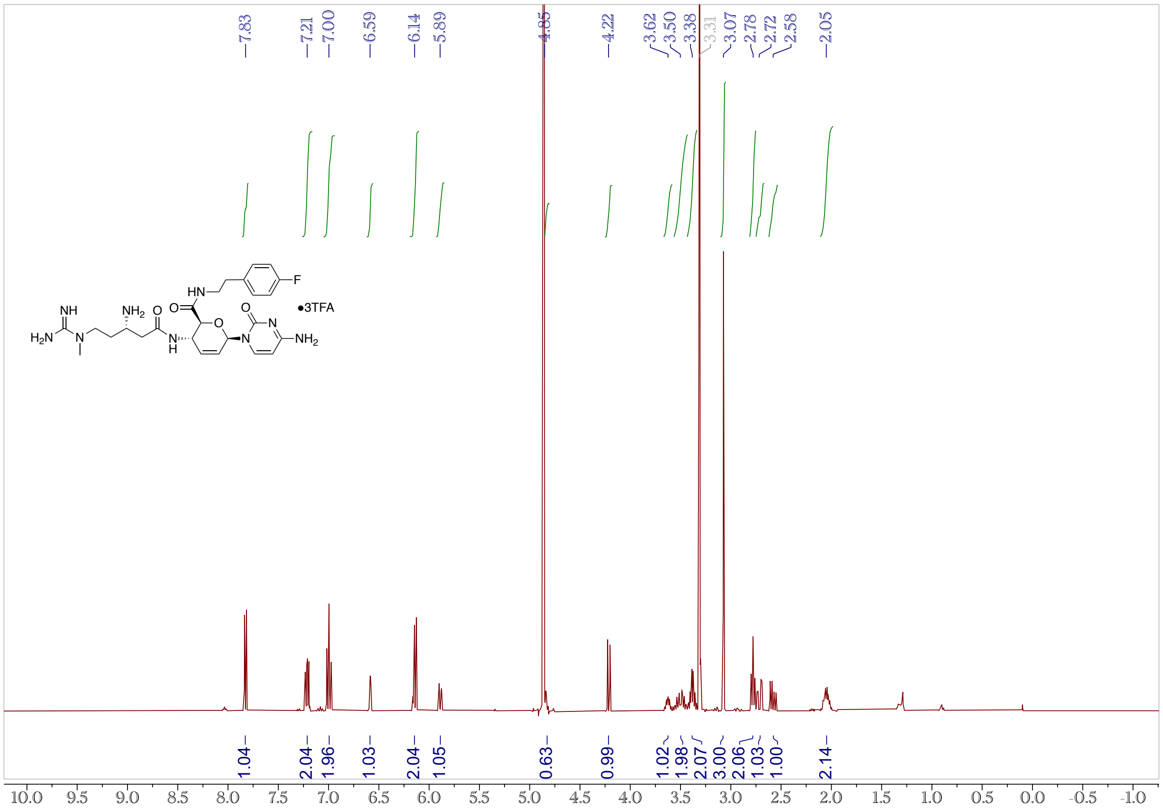


**Figure S12.** $^{13}$C NMR (100 MHz, $CD_3OD$) Spectrum of **5e**.

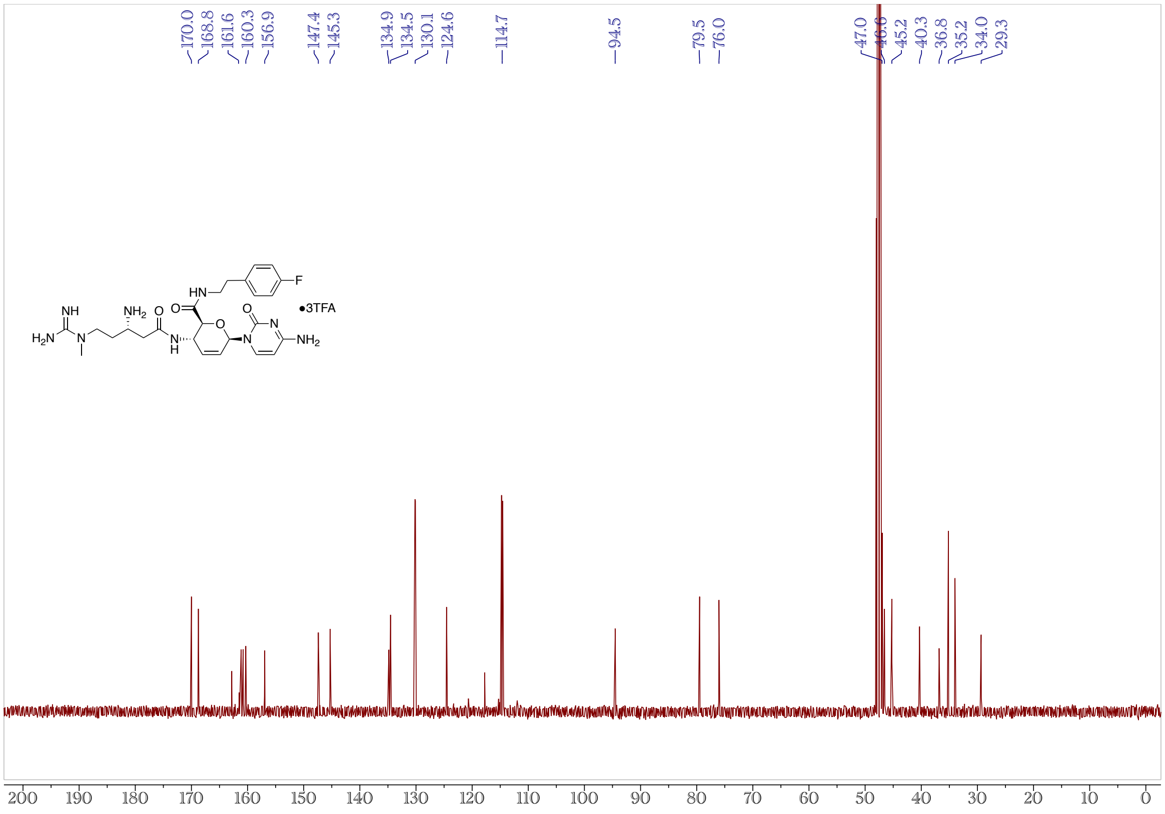

**Figure S13.** $^{1}$H NMR (400 MHz, $CD_3OD$) Spectrum of **5f**.

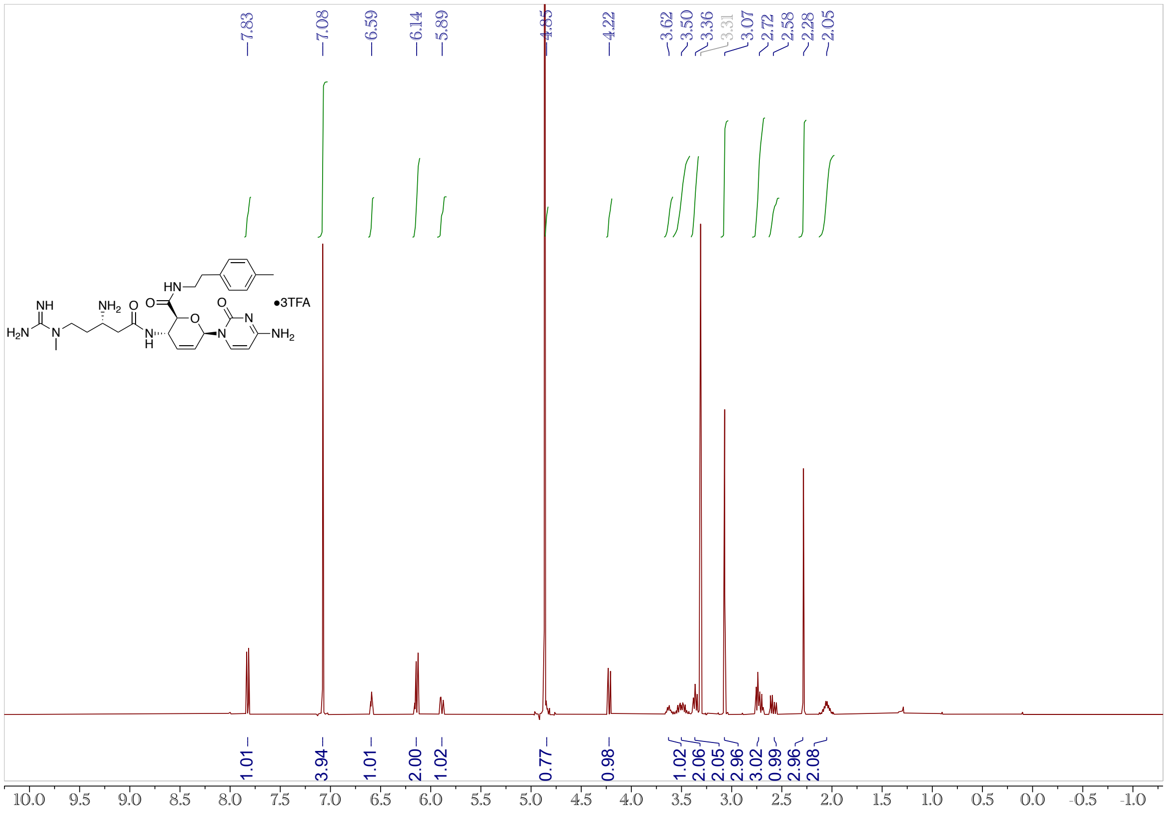


**Figure S14.** $^{13}$C NMR (100 MHz, $CD_3OD$) Spectrum of **5f**.

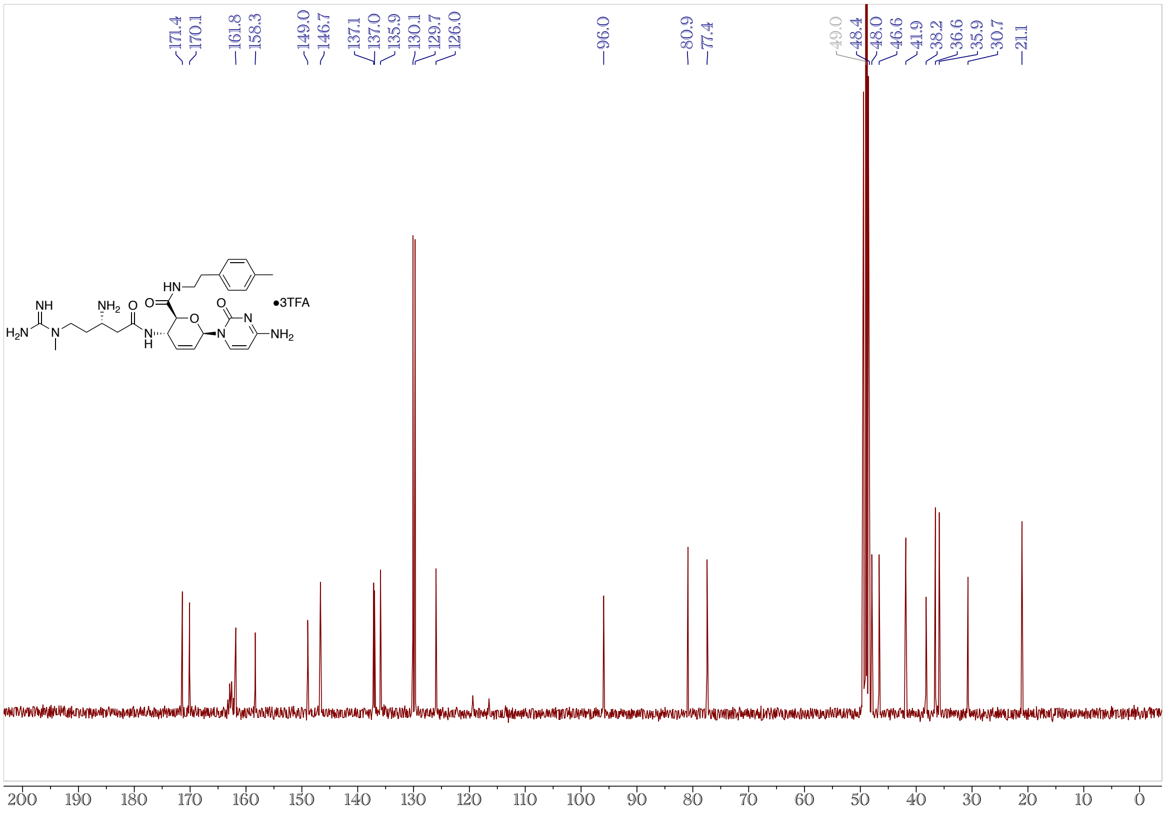

**Figure S15.** $^{1}$H NMR (400 MHz, CD$_3$OD) Spectrum of **5g**.

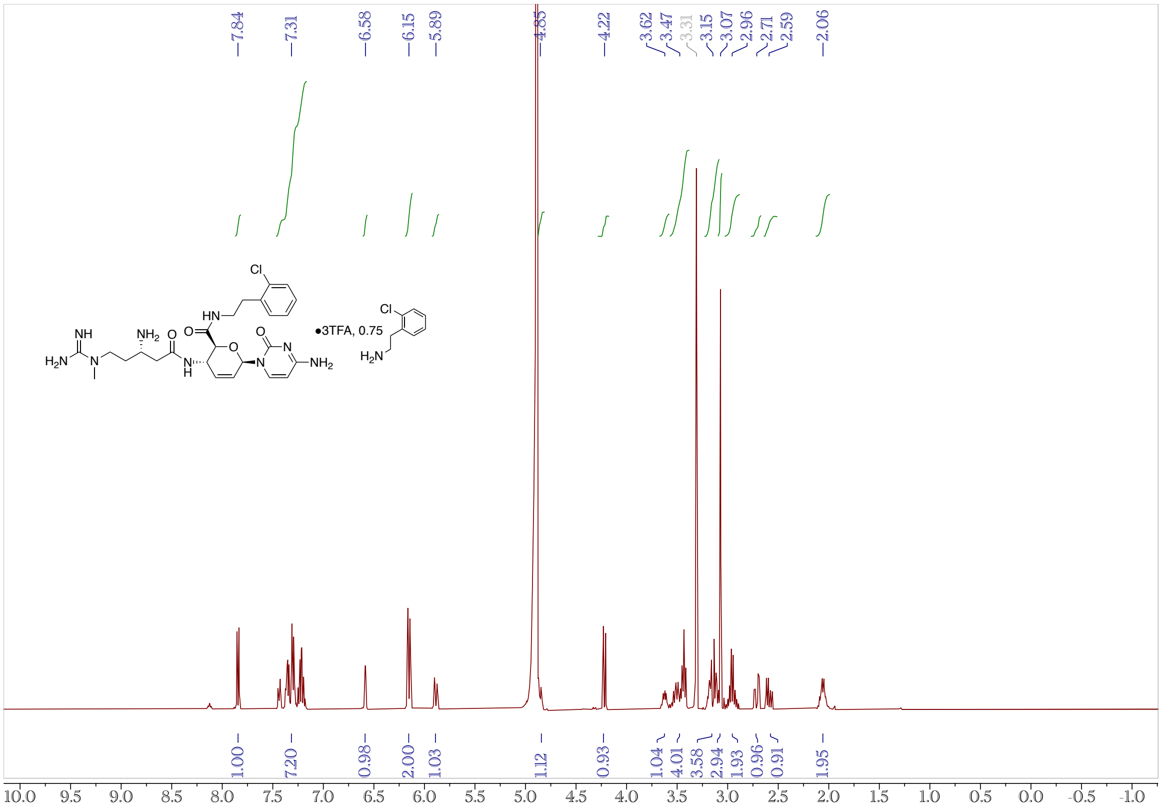

**Figure S16.** $^{13}$C NMR (100 MHz, CD$_3$OD) Spectrum of **5g**.

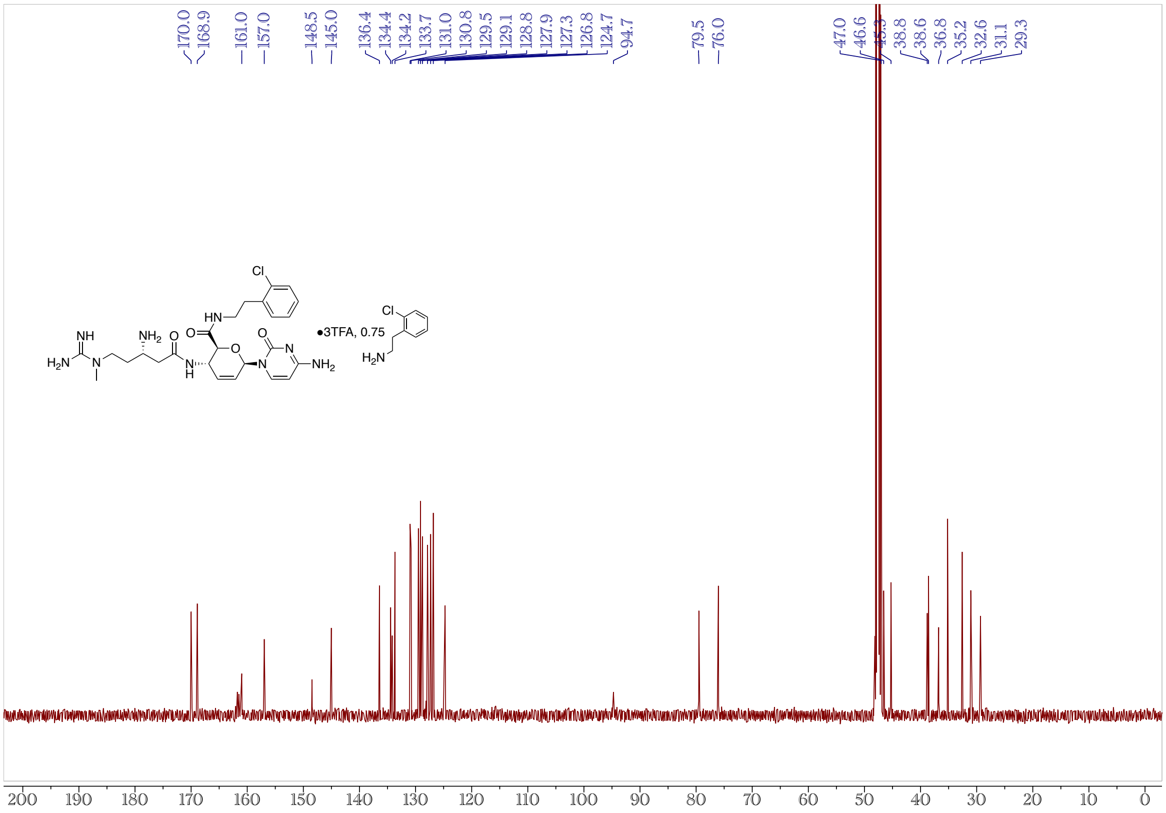

**Figure S17.** $^{1}$H NMR (400 MHz, $CD_3OD$) Spectrum of **5h**.

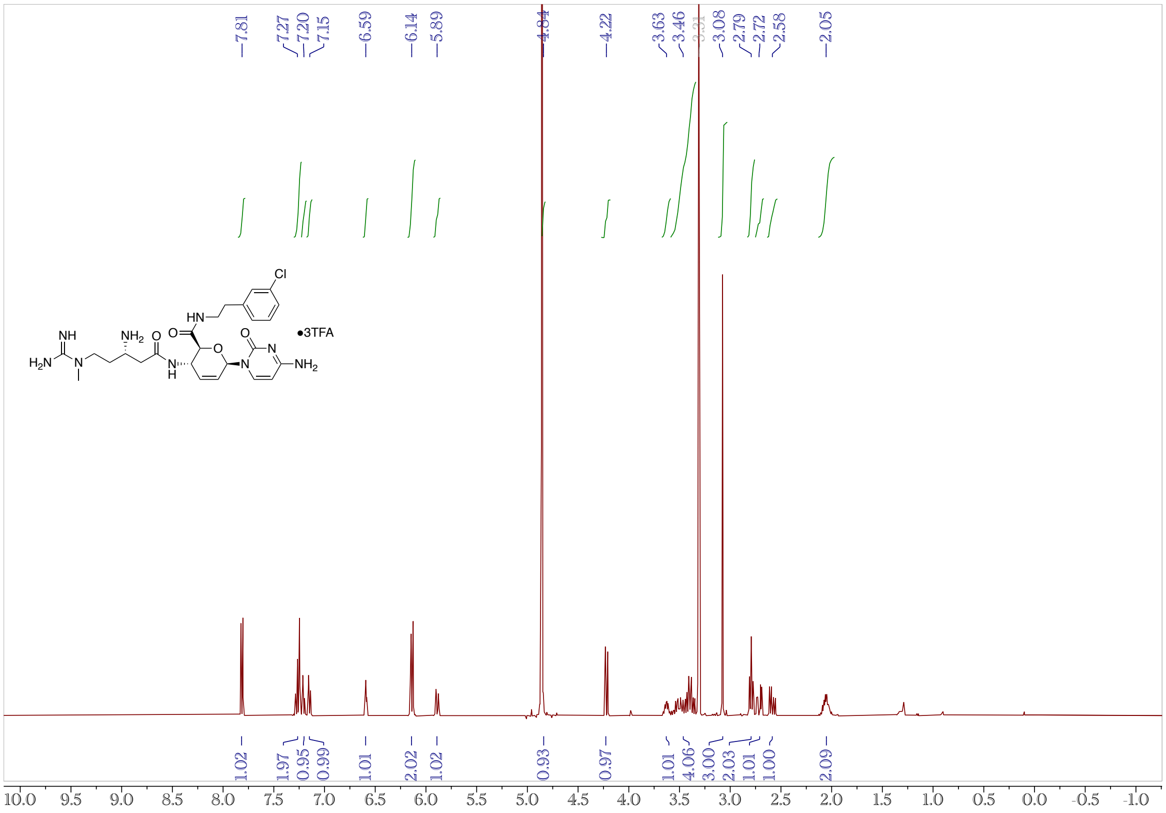

**Figure S18.** $^{13}$C NMR (100 MHz, $CD_3OD$) Spectrum of **5h**.

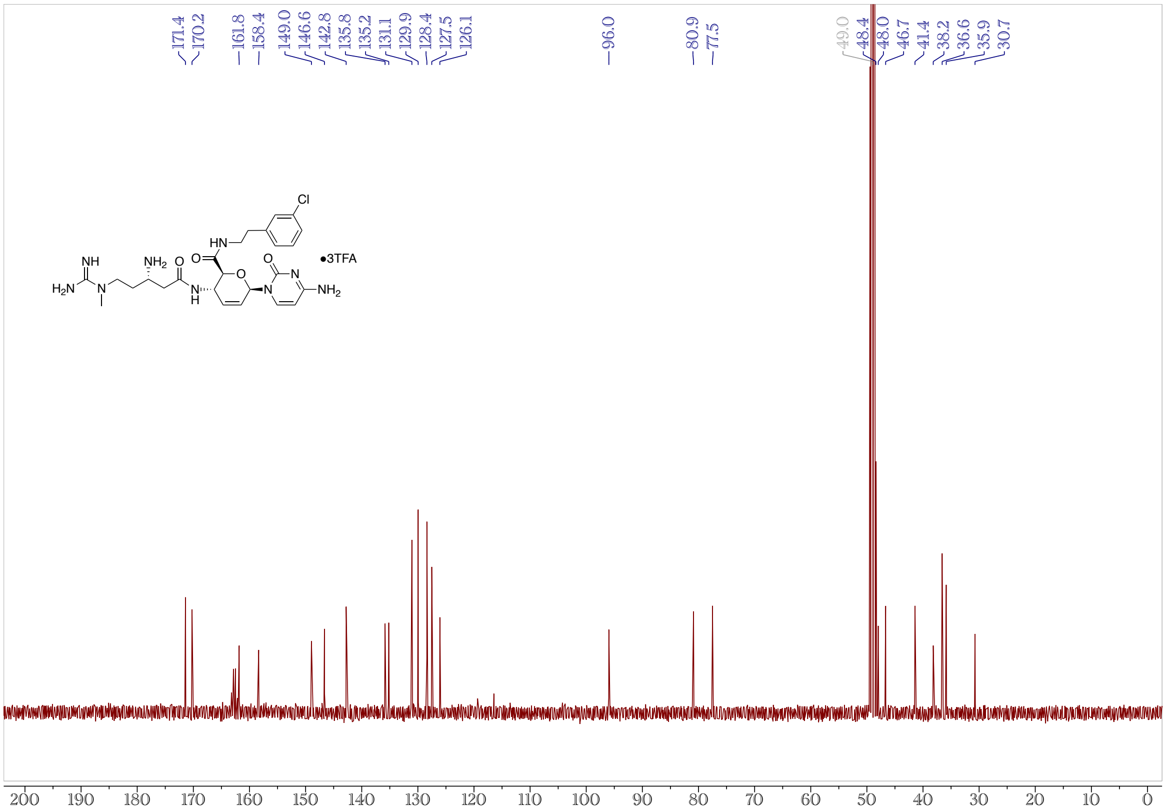

**Figure S19.** $^{1}$H NMR (400 MHz, $CD_3OD$) Spectrum of **5i**.

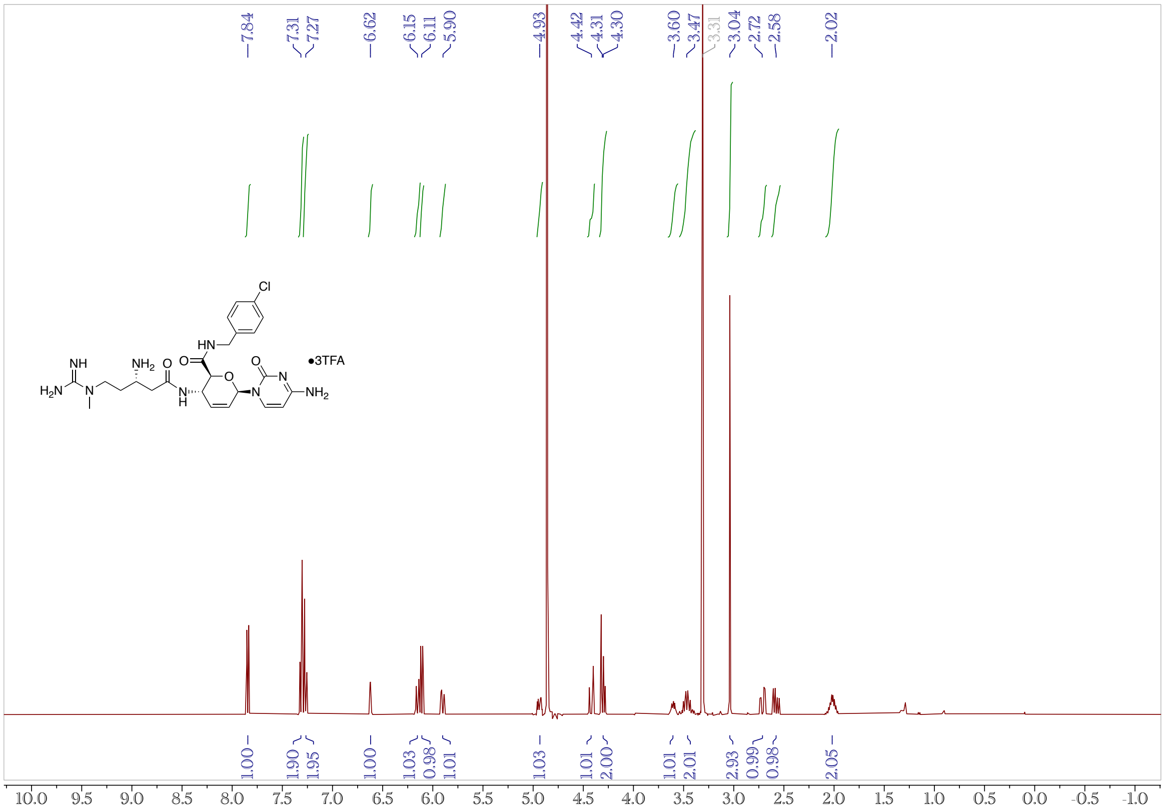


**Figure S20.** $^{13}$C NMR (100 MHz, $CD_3OD$) Spectrum of **5i**.

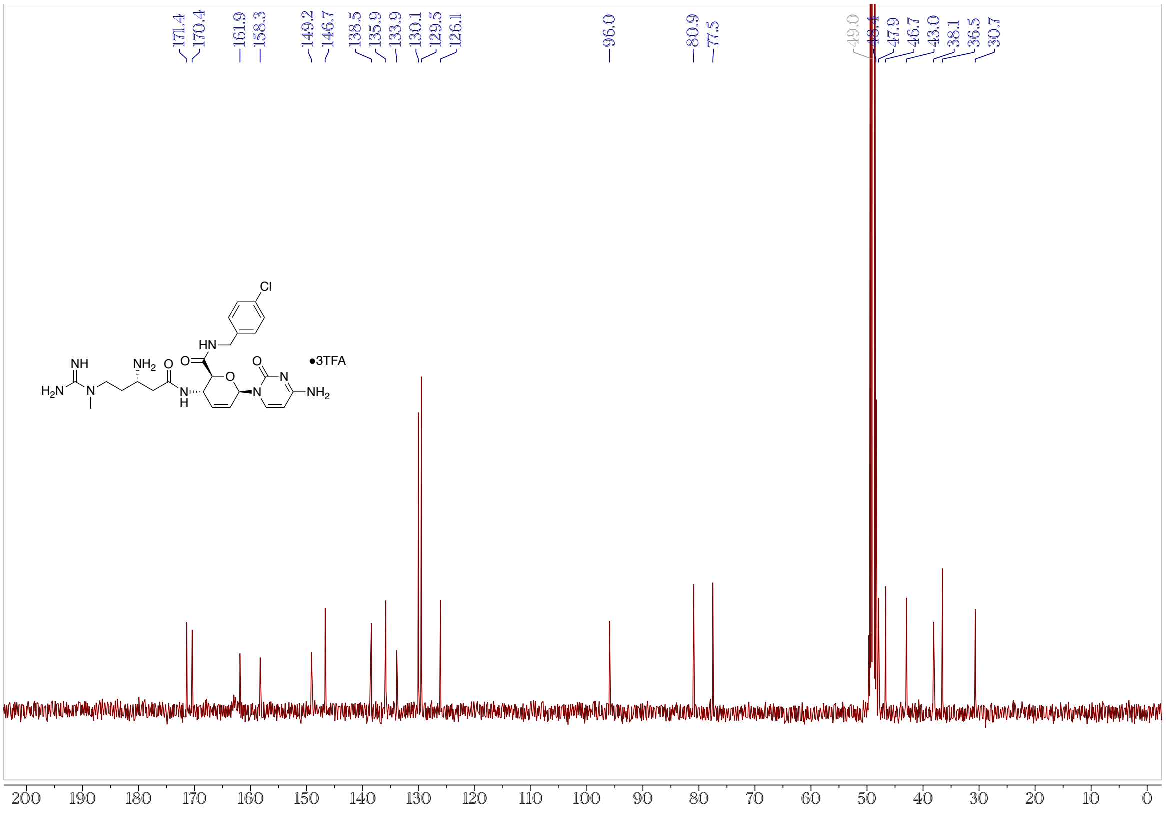

**Figure S21.** $^{1}$H NMR (400 MHz, $CD_3OD$) Spectrum of **5j**.

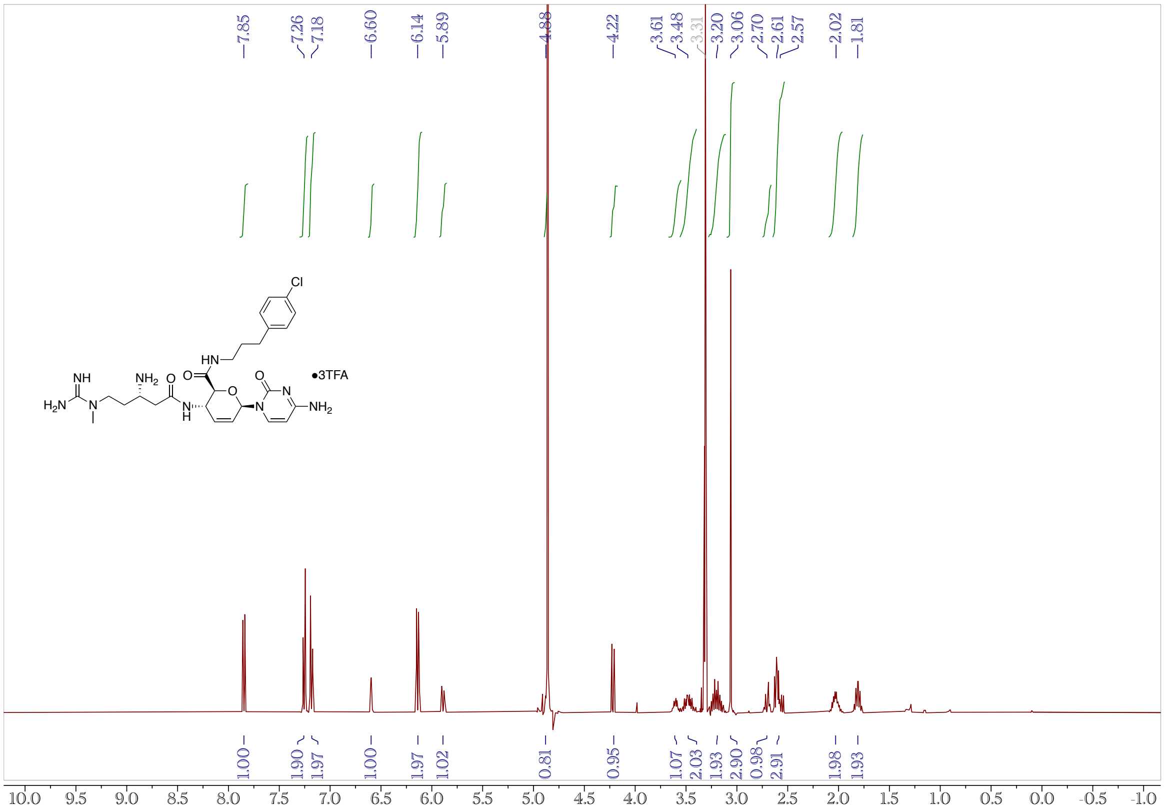

**Figure S22.** $^{13}$C NMR (100 MHz, $CD_3OD$) Spectrum of **5j**.

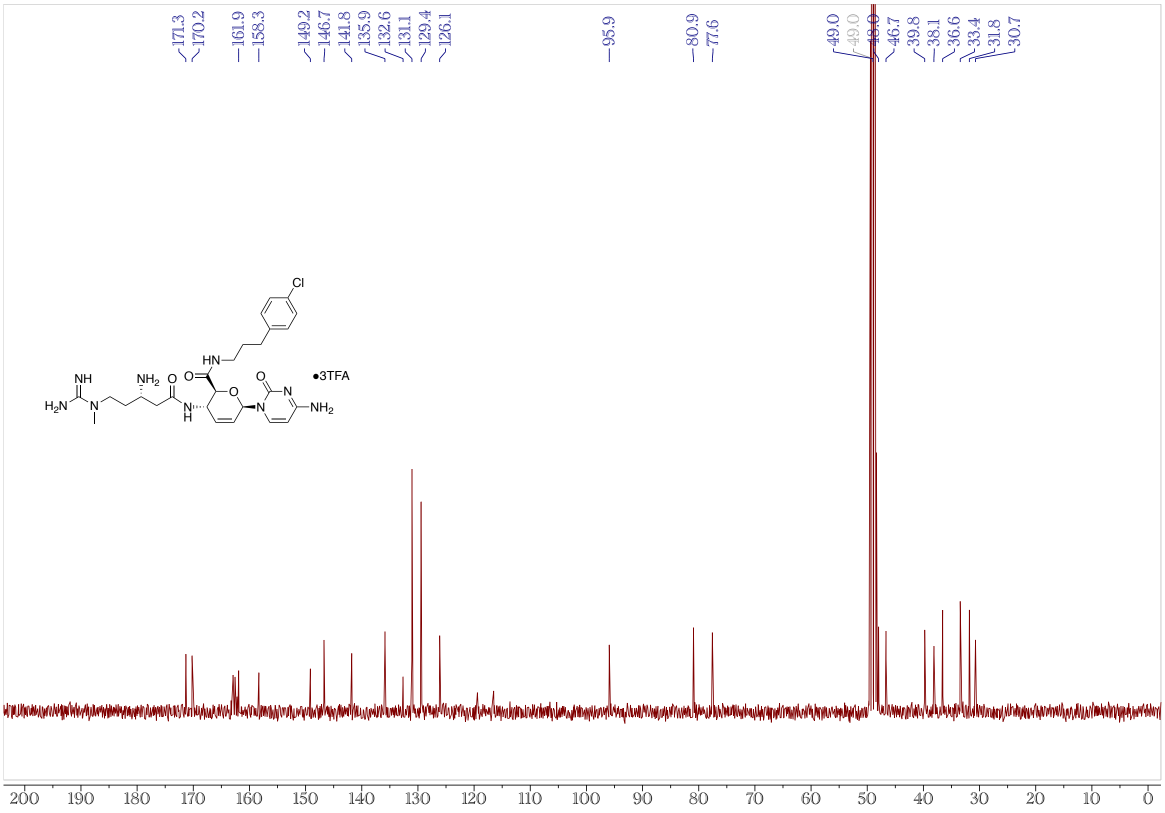
